# Natural Induction: Spontaneous adaptive organisation without natural selection

**DOI:** 10.1101/2024.02.28.582499

**Authors:** Christopher L. Buckley, Tim Lewens, Mike Levin, Beren Millidge, Alec Tschantz, Richard A. Watson

## Abstract

Evolution by natural selection is believed to be the only possible source of spontaneous adaptive organisation in the natural world. This places strict limits on the kinds of systems that can exhibit adaptation spontaneously, i.e. without design. Physical systems can show some properties relevant to adaptation without natural selection or design. 1) The relaxation, or local energy minimisation, of a physical system constitutes a natural form of optimisation insomuch as it finds locally optimal solutions to the frustrated forces acting on it or between its components. 2) When internal structure ‘gives way’ or accommodates to a pattern of forcing on a system this constitutes learning insomuch as it can store, recall and generalise past configurations. Both these effects are quite natural and general, but in themselves insufficient to constitute non-trivial adaptation. However, here we show that the recurrent interaction of physical optimisation and physical learning together results in significant spontaneous adaptive organisation. We call this adaptation by natural induction. The effect occurs in dynamical systems described by a network of viscoelastic connections subject to occasional disturbances. When the internal structure of such a system accommodates slowly across many disturbances and relaxations, it spontaneously learns to preferentially visit solutions of increasingly greater quality (exceptionally low energy). We show that adaptation by natural induction thus produces network organisations that improve problem-solving competency with experience. We note that the conditions for adaptation by natural induction, and its adaptive competency, are different from those of natural selection. We therefore suggest that natural selection is not the only possible source of spontaneous adaptive organisation in the natural world.

## 1. Introduction

Natural selection is a process of adaptation characterised by the differential survival and reproduction of randomly varying types (Lewontin 1970, Futuyma 1979). By describing the mechanism of natural selection, Darwin showed that the number of natural processes that could produce adaptive organisation spontaneously (i.e. without a designer) was at least one - not zero, as previously thought. Since he showed that it is possible for natural processes to produce adaptive complexity spontaneously, are we sure that the number of such processes is *exactly* one? Could there be others? Or would all putative alternatives actually turn-out to be either some form of natural selection in another guise, or not a genuine source of adaptation at all (Dawkins 1983, Watson and Lewens 2024)?

### 1.1. Natural sources of adaptation

If natural selection is understood as a specific type of process (i.e. not defined as any and all processes that produce adaptation), and likewise if adaptation is not defined as the product of natural selection, then it is a valid question to ask whether other processes can produce adaptation. Whilst no well-informed thinker doubts that evolution by natural selection plays a central role in explaining the adaptive organisation of biological organisms, and the role of natural selection in biological evolution need not be threatened by asking this question, the potential ramifications of such a possibility could be important to biological understanding in many areas. These include: understanding how natural selection interacts with other biological and physical processes (e.g. development, niche construction, ecological dynamics, extended inheritance mechanisms, learning, organismic agency) (Laland, Uller et al. 2015, Watson and Thies 2019, Levin 2023); understanding how natural selection got started (Cairns-Smith, Hartman and Cairns-Smith 1986, Nowak and Ohtsuki 2008); informing open questions such as how natural selection rescales from one level of organisation to another, i.e., evolutionary transitions in individuality (Maynard Smith and Szathmary 1997, Watson, Levin and Buckley 2022); and exploring the possibility of adaptation in biological systems that are not evolutionary units (Lovelock and Margulis 1974, Levin 1998, Power, Watson et al. 2015, Wilson 2016). See (Watson and Lewens 2024) for further discussion. So, to explore further the systematic investigation of the question that Darwin first posed, what kinds of natural processes can produce adaptive organisation spontaneously?

Actually, it is clear that other mechanisms of adaptation exist in biology, such as adaptive plasticity and animal learning (West-Eberhard 2003, Ghalambor, McKay et al. 2007), and also in engineered systems, such as machine learning mechanisms (Mitchell and Mitchell 1997). These do not necessarily depend on variation and selection for their operation. For example, adaptation like that which occurs via the organisation and reorganisation of synaptic connections in the brain need not involve a selection process^1^. Machine learning algorithms, likewise, can utilise gradient methods that do not require any variation and selection. Of course, these examples involve specific adaptive mechanisms that are themselves selected or designed for the purpose of producing adaptive outcomes. If this neural machinery, for example, is complex and specific then its origination requires explanation, even if the adaptation that occurs thereafter (after the machinery has been ‘set-up’) does not involve natural selection. Thus, the kind of adaptation that brains (and machine learning systems) demonstrate does show that natural selection is not the only possible adaptive mechanism, but they do not appear to offer a solution to both the chicken problem and the egg problem, namely, to present an adaptive mechanism that is not natural selection and also does not need natural selection to explain how it arose. What adaptive mechanism could be so simple that it does not require selection or design either for its operation (at ‘run time’) nor to explain its necessary machinery (i.e. for construction or ‘set-up’)? For lack of any such known alternative in the natural world, natural selection is understood to be the *source* of *all* adaptive organisation (Dawkins 1983). Moreover, it has been argued, quite convincingly, that natural selection is the only possible mechanism of adaptation that could occur naturally (even on another planet where we could imagine biology that worked differently), hence “universal Darwinism” (Dawkins 1983) (Lewens and Watson 2024, Watson and Lewens 2024). Is it really impossible for there to be other sources of spontaneous adaptation besides natural selection? It is worth noting that an unthinking incredulity – as if we already know that spontaneous adaptation is impossible, unphysical, opposed to the natural order of things or even heretical – was exactly the mindset that Darwin already overcame.

If we wish to explore the possibility of a different source of adaptation, i.e. not requiring natural selection either to set-up the necessary machinery or in its operation thereafter, we had better be able to show how it works in a system that is not associated with natural selection at all, i.e. that does not depend on any specific characteristics of biological systems or materials. Even if our real interest in adaptation were exclusively biological, it would need to be possible to show adaptation in a non-biological, physical system. This is a stricter condition than Darwin applied – evolution by natural selection is a mechanism of adaptation that presupposes properties of biological systems. It depends on self-reproducing entities that exhibit heritable variation in reproductive success. But given these other (natural and artificial) examples of adaptation in learning processes, one way to potentially expand the scope of adaptive processes is to ask: What other kinds of systems can learn, and in particular can physical systems learn? More specifically, can a physical system exhibit learning spontaneously, i.e. without being selected or designed for that purpose?

### 1.2. Physical learning and physical optimisation

In fact, the principles of learning systems can be very simple, and spontaneous “physical learning” can be readily demonstrated in various kinds of physical systems (Stern and Murugan 2022). These include mechanical, material, molecular and chemical systems, as well as electrical (Kim, Gaba et al. 2012, McGregor, Vasas et al. 2012, Stern, Pinson and Murugan 2020, Stern, Hexner et al. 2021, Wright, Onodera et al. , Parsa, Wang et al. 2022, Stern and Murugan 2022, Venkatesan and Williams 2022). Although the details vary, the underlying principle is the differential ‘accommodation’ of system structures to forces acting on them. This might involve natural properties such as differential deformation (yielding, creeping, ageing, dilation or atrophy) of connections, bonds or flows, or re-arrangements of internal structure, in response to local stresses in those structures (Chvykov, Berrueta et al. 2021, Stern, Hexner et al. 2021, Stern and Murugan 2022). When the internal organisation of a system is incompatible with, or stressed by, the pattern of forces acting on the system from the environment (or from its own dynamical behaviour), that organisation is caused to change by those forces (Fig.1.B). Such systems bend, deform or give way in the direction that accommodates to the specific stresses created by that pattern of forcing. This can result in a sort of memory (or engram) of that forcing, that remains in the structural organisation of the system, and can thereby influence the system’s subsequent dynamics in a manner that reconstitutes or ‘recalls’ the past pattern. A beautifully simple example is shown in paper folding, where up-folds or down-folds, become easier as the paper yields to the forcing it has experienced, leaving a memory that can recreate a given input-output relation or classification (Stern, Arinze et al. 2020).

**Figure 1:**
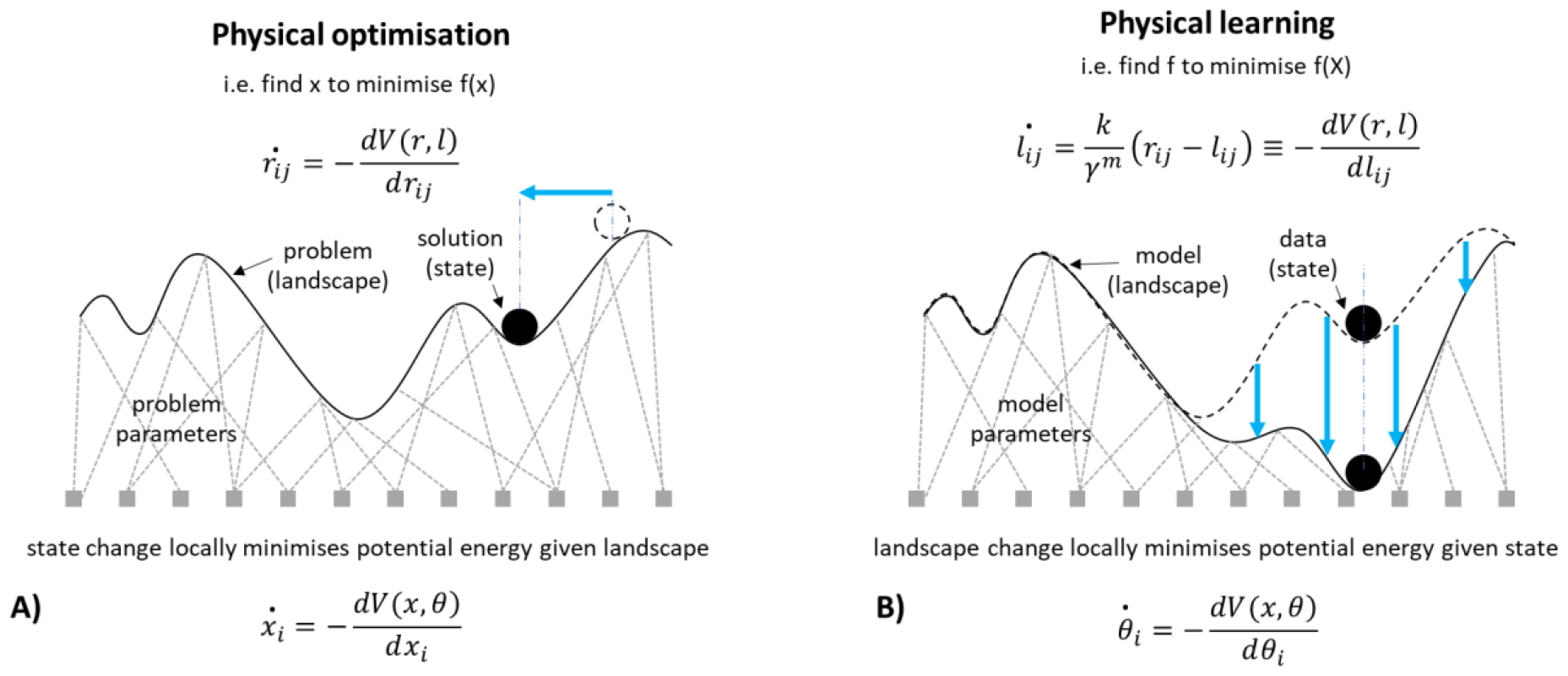
Optimisation and learning in physical systems are complementary processes. A) Physical optimisation is described by the change in state (*x* = {*x*_*i*_}) given the set of parameters (*θ* = {*θ*_*i*_}) defining a potential function, *V*. B) Physical learning, in contrast, is described by the change in the parameters of a model given some state, or distribution of states. For optimisation in masses connected by springs (in an over-damped system), the change in the separation, *r*_*ij*_ , between two masses *i* and *j*, is given by the derivative of potential energy, *V* (a function of all states, *r* = {*r*_*ij*_ }, and all natural lengths, *l* = {*l*_*ij*_ }), wrt *r*_*ij*_ . For learning in masses connected by springs, we show that the change in the natural length of a spring, *l*_*ij*_ , between two masses *i* and *j*, is given by the derivative of potential energy, *V*, wrt *l*_*ij*_ . Thus, in physical optimisation, the states relax to the parameters, and in complement to this, in physical learning, the parameters accommodate to the states. Optimisation is exemplified by a ball that changes position, rolling down a landscape with a fixed shape, and learning is exemplified by a landscape that changes shape, giving way under the weight of the ball with a fixed position. Usually, these two effects are studied in isolation, without feedback.

To make sense of what this means, let’s start simple. In simple forms, a physical memory is trivial. A simple univariate^2^ memory, or imprint, can be shown by something as ordinary as a uniform plastic material like a bed of clay. The features in the pattern of forcing have a one-to-one correspondence with the elements of the material. However, this can only remember one pattern at a time, e.g., the deformation of the clay to a new object overwrites the memory of any previous object. At best, the history of patterns it has been exposed to is reduced to an average or consensus pattern. In contrast, when the deformation in a system occurs in an internal structural organisation that effects linkages or relationships between features, as in these examples, this is capable of a higher-order or associative memory (Watson, Wagner et al. 2014, Watson and Szathmary 2016, Stern, Hexner et al. 2021, Sun, Dong et al. 2021, Stern and Murugan 2022). The deformation in such a system can store and recall multiple patterns with high fidelity (Hopfield 1982, Watson, Wagner et al. 2014). In this case, it is possible for the internal organisation to represent a set or class of patterns, or to learn a non-linear functional relationship between input and output variables. Such physical learning thereby shows competence in some conventional learning tasks such as classification or function learning (Stern, Hexner et al. 2021, Stern and Murugan 2022). Like a simple neural network (which, abstractly, can also be understood as an example of a system with connections that deform under stress^3^), a physical system with this property can also *generalise* to recognise, classify or produce novel patterns that belong to the same general class of patterns even though these specific cases have not been experienced previously.

To understand the potential adaptive properties of physical systems further, let us now begin to relate this kind of learning with energy minimisation and optimisation. Consider the dynamics of a physical system described by the local minimisation of an energy function. Hopfield and Tank (1985) illustrate how this can be interpreted as an optimisation process that can solve optimisation problems. That is, if the interactions between components of the system correspond to the constraints of a problem, then the natural behaviour of those components is to change state in the direction that *relaxes*. That is, in the direction that reduces any stresses caused by those interactions, thereby finding states that resolve the frustrations between state variables caused by violations to those constraints. Thus a physical system, by following the local gradient of its energy function, moves toward better solutions to the constraint problem (Fig.1.A). This proceeds until a local minimum is reached in the energy landscape, corresponding to a locally-optimal solution to the system of problem constraints (Hopfield and Tank 1985).

The physical learning examples described above can now be understood as a special case of this very fundamental energy-minimisation principle. That is, learning can be understood as the local optimisation of model parameters to data, and this can be implemented in a physical system when the internal organisation of a system gives-way under stress. The difference with physical optimisation is that in a simple *optimisation* process we consider how the state of the system accommodates to the landscape of the problem, whereas in a physical *learning* system we consider the reverse, i.e. how the landscape of the model accommodates to the state of the data. Optimisation is a process provided by a dynamical system described by some state variables (solution variables) and some parameters describing the interactions between them (the problem parameters). Learning is a process provided by a dynamical system described by some data and some variables describing the relationships between them (the model parameters). In both cases, the energy function can be the same kind of function – a product of state variables and interaction variables. Learning and optimisation are thus complementary processes (Fig.1); the variables that change in one case are the variables that do not change (the system parameters) in the other. And in both cases, the dynamical behaviour is simply local energy minimisation – either directly in the (ordinary) state variables for an optimisation process, or in the interaction variables for a learning process (Fig. 1).

### 1.3. Induction and deduction

Since learning and optimisation are both described by local gradient-following processes, simply applied to different subsets of variables within a dynamical system, there is a level of abstraction where they can be understood to be examples of the same principle. But it is a mistake to believe there is no conceptual difference between them. A crucial distinction is that optimisation involves a set of state variables (external or observable), and learning involves the *interactions between them* (internal or unobservable). The latter can be understood as *second-order* variables that control the interactions between *ordinary* (first-order) state variables. In Figure 1, the relational aspect of the problem/model is represented by a network of interactions that define the shape of the landscape (shown in the style of Waddington’s ‘epigenetic landscape’ determining the dynamics of the developmental process (Waddington 1957)) (clay, in contrast, might be represented by vertical ties that connect straight down from each point on the surface to a corresponding point on the base).

The conceptual import of this mechanical distinction is that learning systems intrinsically involve *induction*. Induction is the process of inferring general rules from specific examples (Fisher 1935, Solmonoff 1964, Skyrms 1975). For example, *this swan is white, that swan is white → all swans are white*. Unlike deductive inference (which draws specific conclusions from general rules), we immediately notice that inductive conclusions are not logically valid (you cannot conclude with certainty that *all* swans are white if you have not observed all swans). Inductive conclusions are not deductively supported by the evidence – they go beyond the data. Induction is nonetheless necessary for learning with generalisation; the ability to perform well on previously unseen cases or, relatedly, the ability to generate novel examples from the model. Generalisation necessarily requires inferences that go beyond that which is supported by the data (by definition), and without generalisation we have only memory, not genuine learning.

Whilst induction in learning systems can take many forms, the principle is simple^4^. In particular, although the learning process results in a particular model (e.g., “all swans are white”) there may be many models that are equally compatible with the data (e.g. “all animals are white”, “the first two swans are white and all others are pink”, or “swans are always one colour”). In a physical system, this means that there are potentially many energy landscapes that are consistent with the pattern of forcing the system has experienced in the past. This means that its internal organisation is necessarily *underdetermined* by its experience (just as general conclusions are not deductively entailed by specific observations). Nonetheless, when the model deforms, a particular landscape is obtained (rather than *all* the possible landscapes consistent with the data). It thereby represents a particular conclusion that (although consistent with past experience) is neither assured by past experience, and neither is it the only conclusion compatible with past experience. This is why the distinction between ordinary state variables and second-order state variables is important to learning, i.e. because the latter are under-determined by observations on the former, and thereby afford the possibility of generalisation.

Optimisation of state variables on this model landscape (finding a local minimum) constitutes the generative recall of a pattern from the model. This can include the recall of specific states from past experience but also the production of novel instances via generalisation. How the landscape generalises is under-determined by the data and instead depends on the specific nature of the model space (i.e. internal architecture) (Watson 2023). The under-determination of the model and the consequently specific nature of the distribution generated from the model is what makes the optimisation of model parameters in physical *learning* different from the optimisation of (ordinary) external or observable state variables in physical *optimisation*. When a ball rolls down hill on a landscape (physical optimisation), for example, the process is analogous to deduction (not induction); that is, the local energy minimum arrived at, follows certainly from the initial state (specific instance) and the landscape (general rule). Whilst the increments to the model parameters are also determined by the current data point and the current model parameters, the latter are internal variables (general rules) that are not directly controlled by the application of the external forcing (specific instances). These internal variables may depend in principle on the entire history of experiences, and may involve subtle symmetry-breaking dynamics that involve the whole system and dominate the influence of its initial conditions (Watson 2023).

To further clarify, the difference between learning and optimisation tends to zero only if each internal variable has a one-to-one correspondence with an external state variable (like the bed of clay); in this case the model space is degenerate precisely because the model parameters cannot represent any interactions or associative relationships between features of the data. It is this non-relational property that prevents clay from representing underlying structural regularities in the data, and this prevents it from being able to generalise^5^. All the examples of physical learning therefore involve re-arrangements to internal structural variables that are changed by the pattern of forcing applied, but under-determined by the true causal processes that produced the patterns and correlations in that forcing.

### 1.4. What is adaptation?

A system that is deformed or distorted by the forcing applied to it, or simply relaxes under the stresses it experiences, is not (normally) what we mean by adaptation (regardless of whether that deformation involves observables variables or internal structure). Even learning a function or a generalised class of patterns, though it might be analogous to simple forms of cognitive learning, is not the same sense of adaptation that we seem to refer to in biological evolution. Accordingly, the physical optimisation and physical learning examples have not been claimed to be new sources of adaptation (although, learning principles can deepen our understanding of adaptation by natural selection (Valiant 2013, Watson and Szathmary 2016)). So what do we really mean by adaptation? And how does it relate to learning and optimisation?

Definitions of adaptation and adaptive organisation that are tied to natural selection, organismic reproduction or Darwinian fitness, are not useful in answering this question (Watson and Lewens 2024). The ‘appearance of design’ concept (Dawkins 1983) is closer to what we need, because it is not necessarily tied to Darwinian processes, but it is not easily quantifiable. In fact, defining exactly what we mean by adaptation is an open problem in biology. Note that Darwinian fitness, the number of offspring produced, (or inclusive fitness, for that matter), is part of an explanation for biological adaptation, not the explanandum itself (and besides, the relationship between natural selection and maximisation of these quantities is notoriously fraught (Grafen 2009, Birch 2014)). It is not a high number of offspring that needs explaining, and believing that the ‘goodness of fit’ between an organism and environment is *defined* as reproductive output fails to engage with the special properties of the biological organisations that facilitate this. To assess whether there can be any sources of adaptation other than natural selection, it is necessary to let go of the idea that survival and reproduction are the ultimate assessors of adaptation. Whilst these are undeniably important to biological organisms, these are part of the Darwinian explanation for how adaptation happens. In contrast, the goodness of fit between an organism and its environment, and the appearance of design, are much more general notions and more conceptually apt to understanding what adaptation is. However, the former is too easily satisfied in a physical sense (e.g. an imprint in clay) and the latter is rather subjective.

How can we make a general notion of adaptation more objective and quantifiable? In evolutionary computation, the adaptive capability of natural selection is demonstrated through its ability to find good solutions to optimisation problems (Holland 1975). Despite controversy attached to viewing adaptation as a problem solving process, we find it useful to conceptualise adaptation in this way (Fields and Levin 2022)^6^. To formalise this we begin with a notion of searching for points in some configuration space that optimises some quantity, i.e. a process of function optimisation defined over some given space of variables (Grafen 2009, Fields and Levin 2022) (Fig. 1.A). An optimisation ability does not require that a process finds the globally optimal (best possible) configuration (Grafen 2009). This would be too strict: natural selection, for example, does not provide optimal solutions. Conversely, neither do we want to define adaptation in a manner that is trivially satisfied. If we adopt a very simple notion of problem solving, such as that provided by a hill-climbing or local gradient ascent process, that merely finds a local optimum of an objective function, this is trivially satisfied by a physical system as discussed. The behaviour of any physical system that can be described by the local minimisation of an energy function can be interpreted as an optimisation process in the limited sense that it finds locally-optimal solutions to its implicit energy-minimisation ‘problem’ (Hopfield and Tank 1985) (Fig.1.A).

It is not very useful to adopt a definition of adaptation that is satisfied by a literal ball rolling down a literal hill (even if it is functionally equivalent to the process described by natural selection, see Discussion). Instead, we seek a non-trivial problem-solving competency – not necessarily optimal, but not trivial (Watson 2023). Can physical systems do better than the trivial case? Can physical systems exhibit a spontaneous optimisation capability significantly better than a local gradient ascent or descent method? Simulated annealing is a famous example of a computational optimisation process that finds solutions better than local optima (Kirkpatrick, Gelatt Jr and Vecchi 1983), and actual annealing (e.g. in a cooling metal) occurs spontaneously. This shows that the spectrum of possibilities for natural adaptation is non-empty – but still, we aspire to more than cooling metal as a natural example of adaptation. Here we show that the ability of a physical system to *learn* offers the possibility of a physical system that *learns to solve problems better with experience*. Put differently, we classify the ordinary (first order) local energy minimisation behaviour of physical systems as trivial, but optimisation that improves with experience or *second-order optimisation* is algorithmically interesting.

In the previous examples of physical optimisation, the quality of a solution improves over a given state trajectory, but it only reaches a local optimum and its ability to find solutions of good quality does not change with experience. Lower-energy sates, constituting better solutions to the frustrated state variables in the system, may exist but are not obtained. The optimisation ability does not change over time – restarting the system results in the same locally-optimal outcome if it restarts from the same initial position. In the examples of physical learning, a system incrementally improves the fit of an (implicit) internal model to a conventional learning task such as classification or representing a function, e.g. (Stern, Hexner et al. 2021). A problem-solving ability is not demonstrated (except in the same sense of finding a locally optimal fit of model parameters to the data). Both these behaviours (learning and optimisation) are very natural and do not involve particularly limiting assumptions about the system. In both cases the behaviour is determined by local energy minimisation of the same energy function – the only difference is which variables give-way under stress and which are held constant. In one case the state gives way to the problem (optimisation), in the other case, the model gives way to the data (learning) (Fig. 1). But so far, we have considered these two processes in isolation: the outputs of the learning process do not affect the optimisation landscape, and the outputs of the optimisation process does not affect the learning data. This limiting assumption means that learning responds to data that is given by *external conditions* (fixed training data), and optimisation responds to a landscape that is given by *the problem definition* (fixed problem). These effects have been studied separately, but in general dynamical systems both these effects will happen at the same time. What happens to the structure and dynamics of the system in the general case where there is feedback between the two?

### 1.5. Change in state *and* change in interaction structure

What happens when both the state and the structure of a system are variable? If the structure of the system determines the system’s behaviour (i.e. its state dynamics), then changing the structure has the effect of changing the behaviour of the system. But how can a system *change its own behaviour*, let alone *‘improve itself’*? We are specifically interested in cases where the state influences the change in structure and structure influences the change in state through a dynamical feedback. These kinds of systems are studied under other various names. “Meta-dynamical systems” are a general concept to describe systems where the parameters of a dynamical system are, in fact, slow-changing variables (Varela and Bourgine 1992). Examples discussed include evolving gene-regulation networks, neural networks and immune systems. Note that a meta-dynamical system is, from the perspective where all dynamical variables are lumped together, just a dynamical system that *behaves*. But when these variables are separated (into state variables and structural variables), it becomes reasonable to describe it as a system that *changes its behaviour*, i.e. has a behaviour (on a fast timescale) and it changes its own behaviour (on a slow timescale). This is not an arbitrary separation of variables but depends on timescales and also what is observable and what is internal or (not observable) – or, for our purposes, what variables are directly forced and which ones are indirectly forced or induced. In “self-organised” systems, the interaction between state and structure is of various kinds and the emphasis is on the spontaneous organisation of system components, or ‘order for free’ (Kauffman 1993). In “adaptive networks”, the dynamics of a behaviour on a network (e.g. utility-based replication of game strategies) is influenced by the network topology and, reciprocally, changes to the network topology are influenced by the behaviour on the network (Gross and Sayama 2009). This can result in structures that alter state dynamics in predictable ways, e.g. resulting in an increase in equilibrium levels of cooperation (Santos, Pacheco and Lenaerts 2006).

Ashby (Ashby 1952) also studied systems with dynamical state variables as well as dynamical interactions, or wiring parameters, to describe mechanisms of “ultra-stability” and homeostasis. Random re-organisation of the wiring is triggered by stress, until such time as this brings the essential variables back into their viable range. “Adaptive improvisation”, “sequential selection”, and the principle of “least rattling” (Betts and Lenton 2008, Soen, Knafo and Elgart 2015, Chvykov, Berrueta et al. 2021) develop and extend similar ideas; i.e. keep re-organising until an organisation that does not trigger further re-organisation is obtained. These works demonstrate principles similar to those shown here insomuch as there is a dynamical feedback between observed state variables and (internal) interaction variables that control the relationships between the state variables. They demonstrate the ability for such dynamics to satisfice criteria of order and stability but they do not show a second-order optimisation ability, or the ability to find solutions systematically better than local optima.

Since we know that ordinary physical systems can exhibit both optimisation behaviour (given variable state) and learning behaviour (given variable structure), can a system with both flexible state and flexible structure exhibit an increase in problem solving competency spontaneously?

### 1.6. Adaptation by natural induction (a physical model)

Here we examine systems which have the conditions for both physical optimisation and physical learning – systems where both the state and the structure give way under stress, and these changes feedback on each other (Fig. 2). We show that a dynamical system described by a network of viscoelastic connections can spontaneously learn to solve an optimisation problem better with experience under natural conditions. The conditions for this effect are that the connections are viscoelastic, meaning that they give way slightly under stress, and that the state configuration of the system is occasionally disturbed, e.g. subjected to shocks or perturbations. For example, a network of masses connected by springs and subject to disturbances meets these conditions if the springs are not perfectly elastic, i.e. if the springs are slightly plastic, as all physical springs are. The specific self-conditioned organisation of the springs that obtains constitutes an *adaptive* organisation insomuch as it causes the system to generate state configurations that are particularly high-quality solutions to difficult combinatorial optimisation problems. The quality of the solutions discovered can be extremely rare compared to those found by local gradient methods in the same problem, even if such first-order optimisation has repeated attempts. We call the spontaneous feedback between learning and optimisation (occurring without selection or design) *natural induction*, and the ability to improve optimisation ability with experience (i.e. finding solutions better than local optima) *adaptation by natural induction*, to reflect its close association with inductive learning processes^7^ (and to emphasise its complementarity with natural selection).

**Figure 2:**
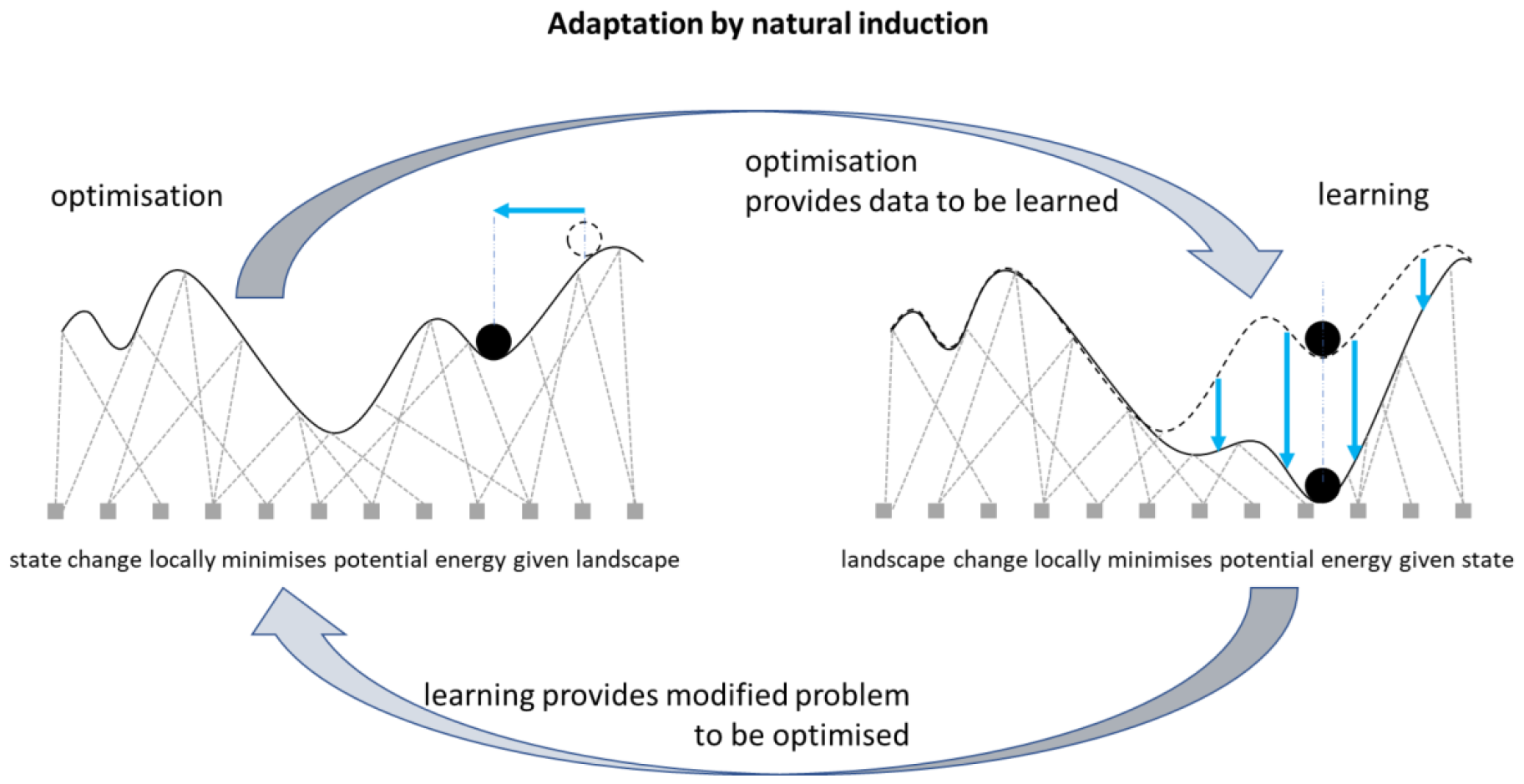
Adaptation by natural induction results from a positive feedback between optimisation and learning. Adaptation by natural induction occurs naturally in physical systems described by a network of viscoelastic connections (subject to disturbances). This involves a dynamical feedback: Energy minimisation on state variables given problem parameters, and energy minimisation on those parameters given the states visited. The initial shape of the problem landscape causes changes in state (optimisation, left). This state at any point in time provides a data point that causes (relatively slow) changes in the problem parameters (learning, right). This provides a modified set of problem parameters to optimise, and so on. Disturbances (randomising the state variables) occur infrequently enough that the system spends most of its time at energy minima (locally optimal solutions) and frequently enough that the deformation of the parameters occurs over a distribution of such optima.

The underlying algorithmic concept is as follows. In each trajectory of an optimisation process, the state changes until it reaches a local equilibrium - corresponding to a locally optimal solution. This optimising process is subject to repeated shocks or perturbations, which effectively randomise the state variables (but not the interaction variables). Each local equilibrium state (achieved in between these disturbances) is thus a point drawn from a distribution of such locally optimal solutions. Meanwhile, the problem parameters give-way slightly in a manner that accommodates to the current state. As the system spends time at each locally-optimal solution visited, the problem parameters model this distribution of state configurations. That is, the problem parameters are model parameters, and the distribution of locally optimal solutions is the ‘training data’. The result is that the system models the outcomes of its own behaviour (hence a “self-modelling” dynamical system (Watson, Buckley and Mills 2011, Zarco and Froese 2018)). In so doing, this changes the dynamics of the optimising system, making it more likely to visit solutions which have already been visited. This changes the distribution of equilibria discovered, which changes the model/problem again, and so on (Fig. 2). This effects a dynamical feedback between learning and optimisation - the state variables give-way in a manner that accommodates to the current system parameters (optimisation), and meanwhile, the system parameters give-way in a manner that accommodates to the current state variables (learning). This results in an increased optimisation ability, in particular in the ability to discover solutions of exceptionally rare quality (Watson, Buckley and Mills 2011). In principle natural induction may occur in any dynamical system where internal structures give-way or re-organise slightly under the stress or forcing a system experiences; the worked example illustrated in this paper uses a network of viscoelastic connections, i.e. springs, which can stand in for many types of network connections in different (biological and non-biological) substrates^8^.

In other settings, the underlying ‘learning to optimise’ principles involved in this effect have been demonstrated in other kinds of self-modelling dynamical systems including neural networks (Watson, Buckley and Mills 2011, Watson, Mills and Buckley 2011, Watson, Mills et al. 2016), gene-regulation networks (Watson, Buckley et al. 2010, Watson, Wagner et al. 2014, Kounios, Clune et al. 2016, Kouvaris, Clune et al. 2017, Brun-Usan, Thies and Watson 2020), social networks (Davies, Watson et al. 2011, Watson, Mills and Buckley 2011), and ecological networks (Power, Watson et al. 2015, Watson, Mills et al. 2016). In all previous cases, however, there was, at some level or another, either a mandated learning process (Watson, Buckley and Mills 2011), a selection process (Watson, Wagner et al. 2014, Power, Watson et al. 2015), or a utility-maximisation process (Davies, Watson et al. 2011, Watson, Mills and Buckley 2011). Uniquely, in this paper we show that the same adaptive principle is exhibited naturally and spontaneously in a physical system – and it is therefore an adaptive process independent of natural selection. Crucially, there is no differential survival or reproduction involved either for the network as a whole, its component parts, or the connections between them; i.e. we demonstrate that natural induction occurs without natural selection (natural selection is neither involved at ‘run time’, i.e. as the adaptation occurs, nor in establishing the initial conditions, the ‘set-up’ or the construction of the system).

In the following experiments we detail example dynamical systems (in two scenarios) each described by a system of masses connected by springs. The first scenario is generic – a network of viscoelastic connections. This illustrates a system which gets better at minimising its own energy function (by changing the organisation of its interactions). The second scenario describes a system that finds good solutions to an independent system of problem parameters (or an external ‘environment’) via changes to its organisation (by changing the coupled dynamics of the system and the environment). This is illustrated for both continuous problem variables (Scenario 2a) and binary problem variables (Scenario 2b). In all three cases, the mechanism of adaptation by natural induction is the same. We present numerical simulations of these systems to illustrate their optimisation capabilities, and demonstrate that they constitute adaptation (by our stringent problem-solving criterion). To conclude, we discuss how natural induction differs from natural selection, and point to some of the important implications.

Our purpose in using a physical model built from masses and springs is not because we are interested in a mechanism depending on these specific physical components, but as an illustration of a general dynamical property in networks of various kinds (biological and non-biological). The purpose of describing this as a system in this physical way is to be as generic as possible and to ensure we are not depending on assumptions that derive from biological systems previously subject to selection. Darwin’s description of evolution by natural selection, though embedded in detailed biological observations (Darwin 1964), was also what we might now describe as an *algorithm*, independent of this biological substrate or any particular instantiation (Lewontin 1970, Holland 1975, Dawkins 1983, Bickhard 2003). In the same way, we aim to use the models that follow as an illustration of an adaptive algorithm that may be instantiated in multiple different substrates (specifically, dynamical systems described by networks of viscoelastic connections, including biological networks of many kinds).

The relevant conceptual territory of this paper thus lies at the interface of biological thinking, physical systems and general dynamical systems. Like the other work in this area (Ashby 1952, Kauffman 1993, Chvykov, Berrueta et al. 2021), we aim to make a small contribution to triangulating the space of possibilities for spontaneous adaptation in natural systems – the aim of this paper is not to displace existing theories but to illustrate additional possibilities and potentialities. This places the work naturally at the cusp of physics and biology (Kauffman 1993), and contributes to understanding their interaction (Salazar-Ciudad, Jernvall and Newman 2003). It is now understood that evolution depends on the interaction of natural selection with the self-organisation properties of non-living matter (Salazar-Ciudad, Jernvall and Newman 2003, Forgacs and Newman 2005, Newman 2022). Natural induction adds to this the possibility that dynamical feedbacks among unorganised components are capable of non-trivial adaptation (not just form and pattern) without the involvement of natural selection.

This concept space, in particular the use of energy landscapes and attractors in biological processes, has long been a part of evolutionary thinking (Waddington 1957, Provine 1989). Unlike much of this prior work, we also use the theoretical framing of computer science and optimisation to provide a rigorous test for non-trivial adaptation (second-order optimiation). This helps to distinguish mechanisms that provide adaptation from other types of complex systems phenomena. Whereas other authors have sought to understand how natural selection might be implemented in primitive physical systems (Cairns-Smith, Hartman and Cairns-Smith 1986), we aim to demonstrate that primitive physical systems correspond to different principles of adaptation familiar in machine learning (Alexander, Cunningham et al. 2021), in particular, adaptation that is *not* natural selection.

## 2. Methods

In the models that follow, we assume that the initial energy function of a dynamical system defines an objective function, i.e. the quality of solutions to a problem (Hopfield and Tank 1985). Initially then, the system finds locally optimal solutions to *its own* ‘energy-minimisation problem’ (Hopfield and Tank 1985). But we define adaptation as doing better than this, i.e. finding solutions better than locally optimal solutions. Logically, this requires that, in order for a physical system to go somewhere different from the nearest locally optimal solution in configuration space, the energy function (describing the dynamics of the system) must be different from the objective function (describing the quality of solutions to the problem) (Watson 2023). In the models that follow, the dynamics of the system changes over time through physical learning, causing the behaviour of the system later in time to deviate from its initial behaviour. To exhibit adaptation, the new behaviour (defined by a new energy function) must cause the system to arrive at configurations that are superior solutions, i.e. better than locally optimal solutions in the original energy function (objective function). A learning system can, for example, accumulate information from experience (i.e. from multiple samples of points in an objective function) about regularities or underlying structure in the problem space, and use this to *optimise better with experience* (Watson, Buckley and Mills 2011). This experience is not provided by an external ‘teacher’, or by any source with privileged information about the problem structure or the location of good solutions (it is ‘unsupervised’ learning, (Watson and Szathmary 2016)). It is acquired through ‘self-modelling’ (Watson, Buckley and Mills 2011), provided by its natural energy minimisation dynamics and the state disturbances, together with the natural generalisation ability of the physical learning process.

We construct a system of *N* masses constrained in a 2D plane (located at *x*_*i*_, *y*_*i*_) connected by Hookean (linear) ideal springs and viscous dampers. Typically, in such systems, pairs of masses are connected by a single spring and damper. With these elements alone the force between each pair of masses is,

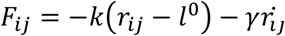

where 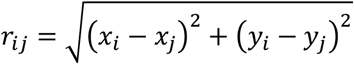 is the distance between the masses, 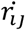 is their relative velocity, *k* is the spring constant, *γ* is a globally defined damping constant and *I*^0^ is the natural length of each spring. In damped systems we expect all the kinetic energy to dissipate until only elastic potential energy remains given by,

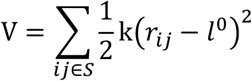

where, {*S*}, is a set of springs. However, real springs are not perfectly elastic, they are imperfectly elastic or slightly plastic, or viscoelastic, meaning that they give way when they are stressed (as is familiar when a spring is stretched too far or held in a stretched position for a long time). A simple example is ‘creep’ where the natural length of a spring increases slightly when it is held under tension or decreases under sustained compression. This slow change in natural length can be modelled by adding a second viscous damper in series with the ideal spring, see Fig. 3.A, which introduces the properties of *Maxwell material* (Roylance 2001).

**Figure 3:**
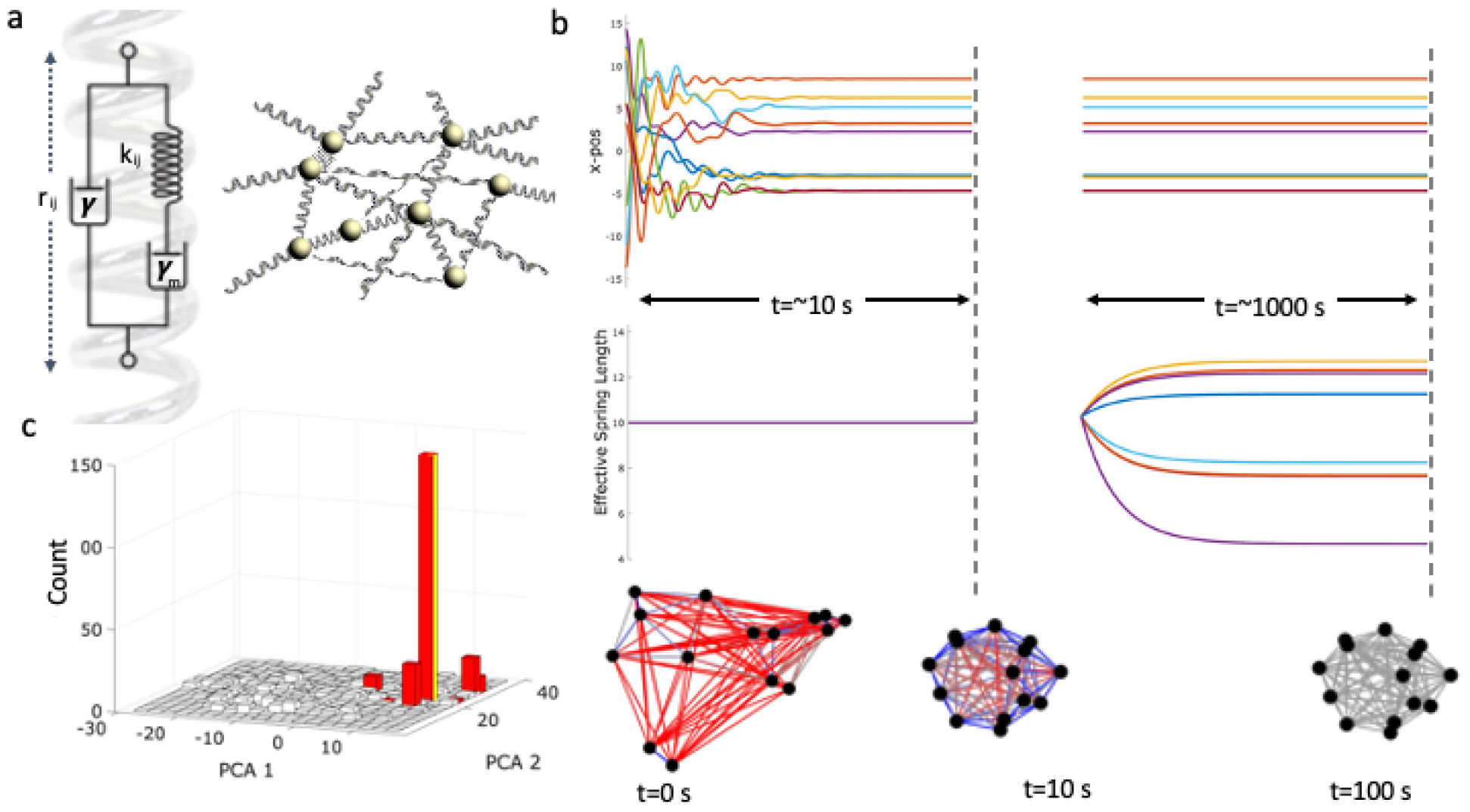
A viscoelastic mass-spring damper system forms a memory of a previously visited configuration. a) Left: A deformable spring is modelled as an ideal spring in series with a viscous damper (γ_m_) which in turn is in parallel with a second viscous damper (γ) (a Maxwell configuration). Right: we simulate a set of frictionless masses in a 2-D plane connected by deformable springs. b) Left: The dynamics of the spring positions in the x-direction settle to stable equilibria over a period of ∼10 seconds (top); there is no appreciable change in the natural lengths of the spring over this period (middle). Right: Over a longer timescale, the positions of the masses are fixed (top) but the natural lengths deform (middle). Bottom row: The network of connected massed in a 2-D plane. Red and blue indicate springs under extension or compression respectively. The initial condition of the masses in the plane (left) over short timescales decays to stable configuration which is under frustration (middle). Over long timescales this frustration is released as the natural length deform (right). c) Histograms of the first two principle components of the pairwise difference between spring positions in the equilibrium configurations reached starting from 1000 initial conditions. Before the deformation of springs, the final configurations are widely distributed (white bins). After the deformation, most initial conditions converged to one of very few configurations (red bins). The most visited configuration aligns with the configuration at which the deformation took place (yellow line). Note: to enable visibility of small values, the tallest red bin is truncated and extends beyond the vertical limit of the plot.

Specifically, in this configuration it can be shown, see Appendix 1, the force between pairs of masses is,

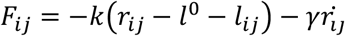

where *l*_*ij*_ is the deformation of the spring and evolves as,

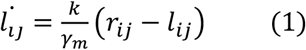

and *γ*_*m*_ is the damping constant of the second spring and we have assumed the timescale of this deformation is slower than viscous drag on spring displacements, *γ*_*m*_ ≪ *γ* (i.e., changes in natural length are slow compared to changes in displacement of the masses). Notice that this deformation of the springs is directional; from Eq. 1 we can see that under tension, or compression, the spring will lengthen or shorten respectively. This deformation described in Eq.1 can also be derived from considering local minimisation energy with respect to spring length, Appendix 1, which concurs with our previous work in Hopfield networks (Watson, Buckley and Mills 2011). Specifically, the potential energy is,

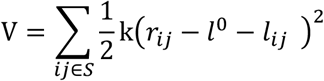

and thus,

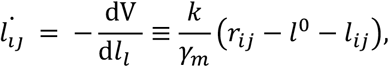

That is, the normal energy-minimisation state dynamics of the system (with perfect springs) describes changes in the positions of the masses given a set of spring parameters, and conversely, the viscoelastic change in the springs is described by the minimisation of *the same energy function* given the positions of the particles. Thus, the particle positions relax – given the frustrations created by the current spring parameters. And, the spring parameters (slowly) relax given the frustration created by the current positions of the particles (Figs. 1 and 2).

As described thus far, the system does nothing very interesting – the fast state dynamics fall to the nearest local optimum, then the slow structural dynamics accommodate to this state, resulting in a new landscape with one memory – at the position of the arbitrary local optimum that was initially found. This therefore affords nothing more than local optimisation (with memory). But the system comes alive when we introduce a ‘pulse’ – repeated disturbances applied to the state variables. This makes the feedback between physical optimisation and physical learning much more interesting. The disturbances effect a sort of ebb and flow, or push and pull dynamic; i.e. the weights push the states, then the states push the weights, and so on – as per the two directions of influence (arcing arrows) in Fig 2. This does not require a mechanism to explicitly interleave these two directions; it is achieved simply by letting the weights change slowly all the time and subjecting the states to occasional disturbances. When the states are in a high-energy configuration they are pushed around by the weights until they reach a local equilibrium and are unable to change any further, and then, whilst they are resting at a local optimum for a long time, the states slowly push on the weights, and then the next disturbance, and so on. Thus, the two directions of influence are implicitly taking turns and the result of the feedback is not simply a single equilibrium state but an organised change in the distribution of a dynamical behaviour.

To the basic setup we introduce occasional disturbances to the state variables (particle positions, not spring parameters). We use three scenarios that differ in the initial arrangement of the system (the ways that particles can move and the arrangement of springs connecting them) to illustrate adaptation by natural induction applied to: 1) solving an intrinsic energy minimisation problem, 2) solving an external optimisation problem (over continuous and discrete variables, 2a and 2b, respectively).

## 3. Experiments and Results

### 1.7. Adaptation by natural induction – generic case

We start by demonstrating the dynamics of a small system and then the ability of such a system to retain information about the configuration of past states. For illustrative purposes we initialise small system with *N* = 15 masses and connect pairs of masses with springs with probability *p*_*c*_ = 90% with uniform natural length, *l*^0^ = 10, and uniform spring constants, *k* = 10.

Over relatively short timescale ( ≈ 10 *s* ) the masses transiently oscillate and settle to an equilibrium, Fig. 3b (upper left). However, note that at this equilibrium not all the springs achieve their natural lengths, and some springs are under compressive or extensive forces, Fig.3.b (bottom middle network, blue and red respectively), and the system is frustrated. Like a neural network with symmetric connections, a physical system of masses connected by springs has only fixed-point attractors (the local minima of the energy function) (Hopfield 1982). In general, such a system may have many such attractors, attaining a variety of equilibria with different amounts of frustration (total energy at equilibrium, *V*).

Over the much longer timescales (≈ 100*s*) the spring lengths change slowly, see Fig. 3.b (right), and the effective natural length, 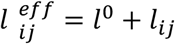 , of each spring deforms in the direction that reduces this tension, see Fig. 3.b (bottom right network). Since the spring lengths are now different, the state dynamics will be altered. But in what way? How will the attractors of the system with the new spring values relate to those of the original system?

To examine how the plastic deformation of springs has affected the stable configurations of the system we plot the distribution of final equilibria (sampled from the same distribution of random starting points) before and after the springs deform. In Fig. 3.c we plot a histogram of the equilibrium states visited from different random initial conditions. To plot this in two dimensions we use the first two principle components of the distances between pairs of masses at equilibrium. The equilibrium reached in any one trajectory depends on the initial conditions (i.e. the starting point of the trajectory). Before the springs deform, we observe a broad distribution of stable equilibrium configurations, Fig. 3.c (blue distribution). One such end point, chosen arbitrarily, is indicated by the yellow column. The system is allowed to rest at this configuration whilst the springs slowly deform. After the springs deform, the system has a new energy function, and its trajectories and equilibria are different (even when using the same initial conditions). We see that the distribution is much more narrowly concentrated, Fig. 3.c (red distribution) and from most initial conditions (> 98%) the masses converge to the yellow equilibrium, i.e. the configuration the system was in when the springs deformed.

Note that this is not simply a system finding stability (in particles and springs), but a system changing its dynamics (by altering its slow variables) in such a way that it can recreate a particular pattern (of its fast variables). This is explained by noting that the effect of spring deformation at any particular state is twofold: it lowers the energy of this particular state (because the springs accommodate to this state and thus resist it less), but more importantly, it increases the size of the basin of attraction for this state (i.e. increases the number of random initial conditions for trajectories that arrive there). This is functionally analogous to the formation of a memory in, for example, a Hopfield network under Hebbian adjustment to connections (Hopfield 1982, Hopfield and Tank 1986, Watson, Buckley and Mills 2011). The changes to the springs thus change the relative size of the attractor basins in the energy function (not just their depth) such that this configuration is visited much more often than others. We interpret this as a memory of a past state configuration – or we might say that the slow system variables have taken an associative ‘imprint’ of its own past state (hence a ‘self-modelling’ dynamical system).

We now examine what happens when springs deform over a distribution of equilibria rather than a single arbitrary equilibrium. The sampling of random initial conditions is provided by the random disturbances; i.e. shocks or perturbations that randomise the positions of the particles. If a system is subject to disturbances of this kind, with intervals of relaxation in between, then the system will visit many different equilibria in proportion to the size of their attractor basins. We assume the time for the fast variables to reach equilibrium is much less than the interval between disturbances, which in turn is much less than the time for the slow variables (spring lengths) to reach equilibrium. Under these conditions, the system spends most of its time at local minima in the energy function but visits many such equilibria on the timescale where the springs deform.

The effect of this is interesting when the number of different equilibria in the initial system dynamics is large. We simulate a system of *N* = 300 with lower connectivity *p*_*c*_ = 50%, with periodic disturbances, see Fig. 4.a. Since the system spends most of its time at equilibrium configurations (not on the transients), the computation is approximated by running the system until it settles to equilibrium and then updating the spring lengths once for each equilibrium visited (rather than updating them at every time step on the transients as well). This is a good approximation in the limit of a slow deformation rate, with sufficiently long periods spent close to equilibrium.

**Figure 4:**
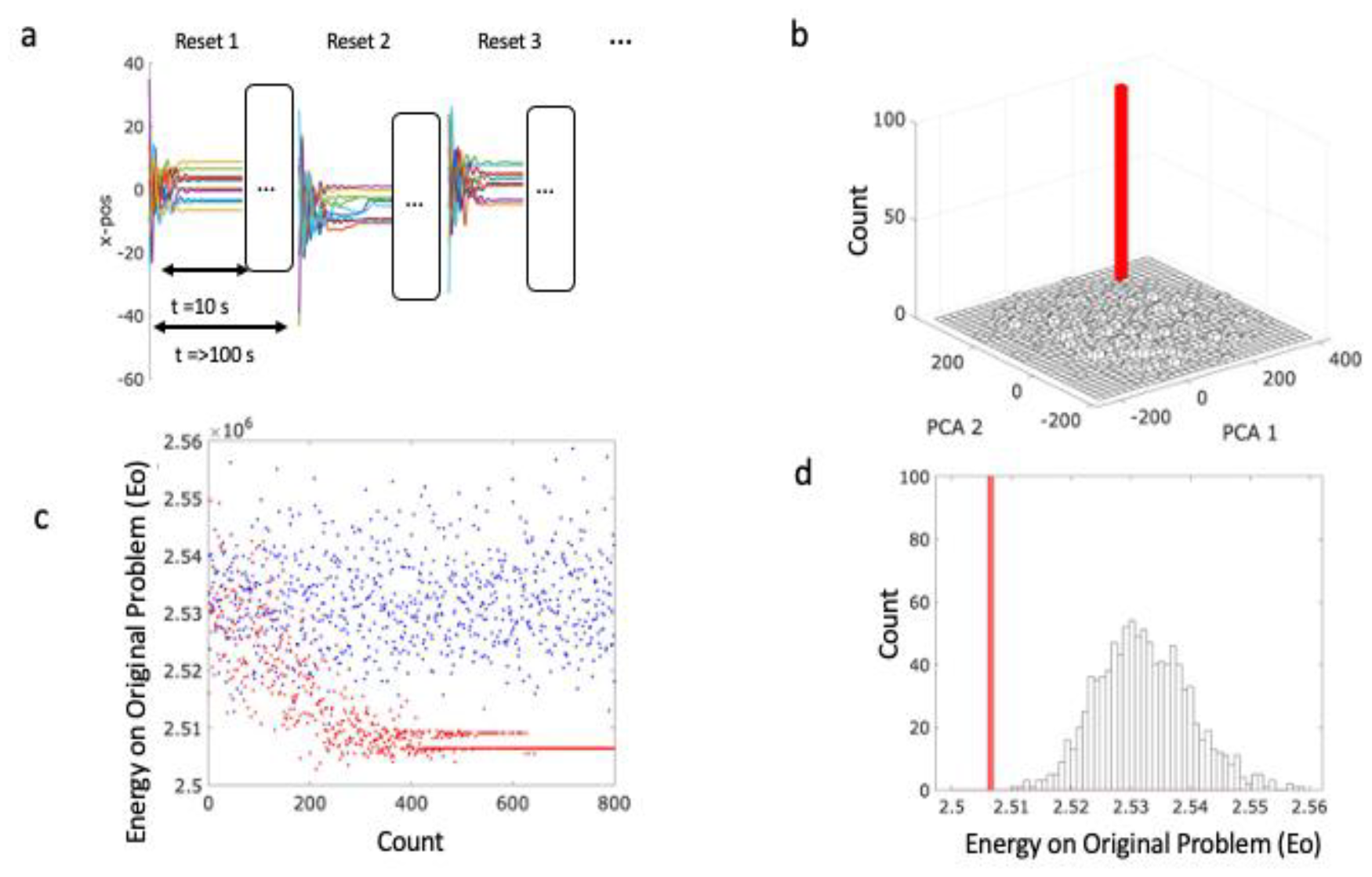
Adaptation by natural induction discovers exceptionally low-energy configurations (Scenario 1). a) The protocol described in terms of the x-displacement of springs over multiple disturbance (resets). The system is disturbed every ∼1000 steps and settles to a stable configuration after which the springs deform. b) Histograms of the principal components of the pairwise difference between spring positions in the equilibrium configurations reached, starting from 1000 random initial states. Before deformation there is a broad distribution of final configurations (white bins), after the protocol in a) all initial conditions converge to single stable configuration (note: the tallest red bin is truncated and extends beyond the figure). c) After each reset we rerun the original spring system initialised at the final learned configuration. The plot shows E_o_ verses the number resets. Without deformation, E_o_ is widely distributed but after many resets, the system consistently finds the basin of attraction of low E_o_. d) The distribution of E_o_ before (white) and after (red).

Although the springs are potentially forming a memory of many state configurations, we find the dynamics of the system with the deformed springs consistently settles to one of very few equilibria as before (Fig. 4.b). This is expected because the system is forming a memory of its own behaviour with positive feedback – the more it visits a state, the more that state is memorised, and the more that state is sampled in future, and so on. But is there anything special about the particular equilibrium (or small set of equilibria) it converges to? Are they good quality solutions, or arbitrary configurations?

As before, the deformation of the springs has the effect of lowering the energy of state configurations it has visited previously, and widens their basins, making them more likely that it visits them again. To assess their quality as solutions to the original energy minimisation problem, we need to examine the configurations found with the new system dynamics and report the energy that these configurations had in the *original* system dynamics. Since the springs now have different lengths, it is possible that the new attractor states may not have been attractor states in the original system. So instead we find which attractor (of the original system) it is closest to, or more exactly, which equilibrium state it is attracted to. That is, we use the final state configuration found in the new system (with the new energy function) to set the initial conditions of the original system and let this system settle (in the original energy function). (For the purposes of taking these statistics, the spring lengths are unaffected by this side assay).

We plot the equilibrium energy of the original system (V_o_) obtained by this procedure over time, as measured by the number of disturbances, thus tracking how this energy changes as the springs deform, see Fig. 4.c (see blue dots). For reference we also plot the energy of equilibria in the original system when starting from the same initial conditions i.e. caused by identical disturbances see Fig. 4.c (red dots). We observe that as the springs deform the system is caused to find lower and lower energy configurations, finally converging on a small number of very low energy configurations see Fig. 4.c (blue dots). In Fig. 4.d we plot a histogram of stable equilibria found from random initial conditions before and after spring deformation (resettled with the original spring values), blue and red histograms respectively. We note that the new distribution is converged on states that are particularly low energy, lower than any of those found in the original behaviour of the system (from the same distribution of initial conditions).

We stress that this is not because it is finding the same arbitrary configuration and lowering its energy, but because it is finding different configurations. These low-energy configurations were present as attractors in the original system dynamics but their basins of attraction were very small and were thus very rarely found with the original dynamics (indeed, not found at all in the number of samples shown). The new system dynamics has enlarged the attractor basins for these particularly good quality solutions, even though we would not even expect them to be visited, even once, under the original dynamics with this number of samples. How is this possible?

The explanation has two parts. First, in systems built from a large number of pairwise interactions, the lower-energy equilibria tend to have larger basins of attraction (by virtue of limits on the slope of the energy function arising from being a sum of many forces) (Watson, Buckley and Mills 2011). It is thus expected that the attractors that are better quality solutions to the energy minimisation problem are visited more often than low-quality solutions (this is essentially why gradient methods, like simulated annealing, have some success in general energy landscapes/objective functions). With the positive feedback on the system dynamics provided by spring deformation, we might expect the system to converge on the largest attractors and these tend to be the best attractors. By virtue of the fact that this makes a system visit good quality solutions more reliably over time, this is already a type of adaptation by our strict criteria, i.e., a system that learns to optimise better with experience. But this simple kind of reinforcement learning is not the whole explanation and the results are significantly more interesting than this reasoning suggests. Specifically, the system finds solutions that are better than any solution found by the original dynamics. This means that the system is not just forming a memory of low-energy configurations it had already visited, it is visiting configurations that are novel (and even lower in energy). This is possible because an associative memory can *generalise* – it can generate novel patterns from the same class, not just patterns it has been trained on (i.e. already visited) (Watson, Buckley and Mills 2011). This is possible because the new spring organisations are not just a memory of past states but an induced model, that goes beyond the training data, to generate new configurations with similar structural regularities.

To quantify this, and explore the robustness of the result to different realisations, we quantify how unlikely the learned energy was to be found in the unlearned system as the number standard deviations (STD’s) between the mean energy of equilibria found in the original system and the minimum final learnt energy. Over 10 runs, we found that 80% had more than 3 STDs from the mean and the median number was 3.4 STDs (we calculated this on the log energy which is better represented by a normal distribution).

Note that if the original spring parameters represent the constraints of an optimisation problem, then the original state dynamics just find locally optimal solutions to that problem (in proportion to how likely they are to be sampled) (Hopfield and Tank 1985). In contrast, the state dynamics of the new system, deformed under these conditions, finds configurations that are much better than any of the locally optimal solutions to the problem found in the same number of samples from the same initial conditions. In other words, through a natural stress-reduction process arising spontaneously from the deformation of the springs, the system learns to solve the problem better with experience.

### 1.8. Solving ‘external’ problems by natural induction

In Scenario 1, the effect of adaptation by natural induction is demonstrated under general and natural conditions, i.e. for any network of viscoelastic connections. Scenario 2 examines cases that demonstrate the same effect under different conditions, changing the way the problem is encoded into the system and the way that a solution is ‘read off’. In all cases, the mechanism of adaptation by natural induction that finds the solutions works in the same way.

In Scenario 1, the springs that represent the problem and the springs that deform are the same springs (we compare the system behaviour before and after deformation by using a copy of the original springs). It is a system which gets better at minimising its own energy function by changing the organisation of its interactions^9^. Can natural induction solve a problem that is external to the learning network? That is, a scenario more analogous to the conventional (though artificial) separation of organism and environment? In this second scenario, we examine a system that finds good solutions to an independent system of problem parameters or an external ‘environment’, via changes to its organisation (by changing the coupled dynamics of the system and the environment).

In Scenarios 2 and 3, the system has two different types of springs – springs that are deformable and springs that are not. The non-plastic springs represent the problem constraints (or external environment), and the springs that are plastic effect the inductive learning (the adaptive system). Thus, the plastic springs modify the total system dynamics in the same way as before, finding configurations that solve the constraints defined by the problem springs. Eventually the plastic springs ‘melt away’ completely, leaving only the unchanged problem springs to determine the final attractor state. In this way, this scenario uses two types of springs and demonstrates the same increase in problem-solving competence without needing to do the side-assay where we transferred the new state back to the old spring values and re-settled the system. This set-up also offers an interpretation where the problem springs represent a problem external to the system (in the environment) and the deformable springs represent an adaptive system (organism or agent). These are coupled together through the shared state variables (an interface or phenotype) to produce a combined dynamics (organism and environment in interaction). The effect of connecting these two networks together is that the environmental dynamics induces a model of itself into the agent’s internal structure (the deformable springs), and the generalised nature of this model’s dynamics has the effect of steering or chaperoning the interface variables into increasingly superior solution states (until leaving them in a solution state as their influence attenuates).

To model this we introduce a second type of spring and distinguish between problem springs (P-springs) and learning springs (L-springs). Specifically, P-springs are perfectly elastic and do not deform, i.e, see Fig.5.a, left. These springs define the energy landscape of the *problem* to be solved, i.e., we interpret the problem as one to find the lowest energy configuration of the P-springs. The L-springs have the Maxwell configuration previously described, see Fig.5.a, right. These springs are highly connected (almost fully), weaker and uniform in natural length and spring constant (as before). The intuition is that the energy landscape of the initial system (with P-springs and initial values of L-springs) will be still dominated by the stronger P-springs. To confirm this we plot histograms of the energies of equilibria found when just P-springs are present against and those found by first running on both P-springs plus L-springs but then removing L-springs and letting it resettle (as per the previous assay). These histograms were almost identical, see Appendix 2. We can thus think of L-springs as the addition of an initially neutral field of plastic material that (initially!) does not alter the problem or the physical optimisation dynamics, and eventually will melt-away to nothing leaving only the bare problem. But in between these beginning and ending time points, the influence of the L-springs does something very interesting to the system’s final resting position.

**Figure 5:**
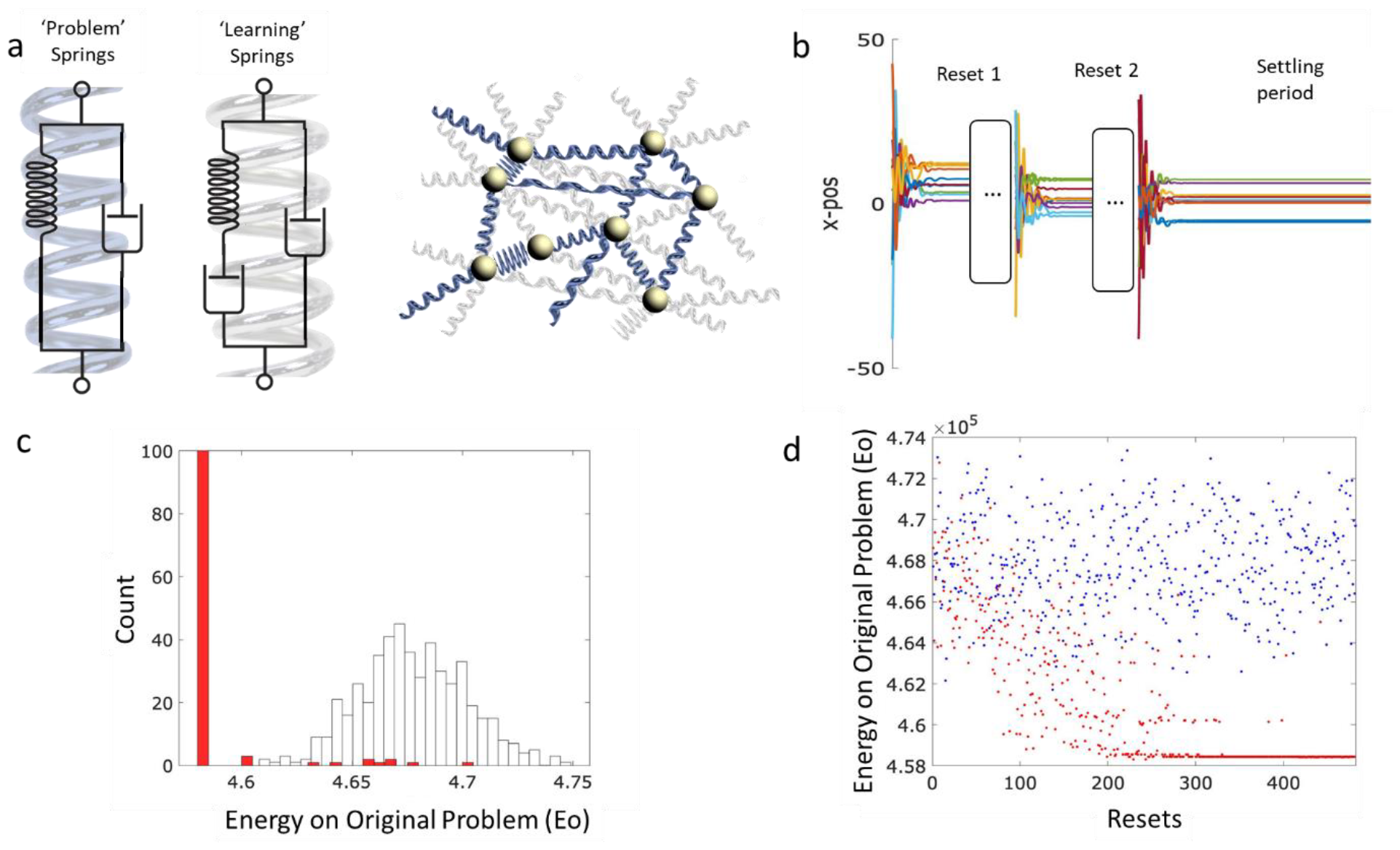
A viscoelastic network discovers low energy configurations of a network of ‘problem’ springs (Scenario 2). a) A material comprising a set of highly connected viscoelastic springs (left most spring configuration and blue) and a relatively sparse set of the perfectly elastic springs (middle spring configuration and grey). b) The same protocol described in Fig. 2 but the system is allowed to settle after disturbances stop. After this period all tension is released in the viscoelastic springs and they no longer contribute to the energy. c) The distribution of energies starting form random initial conditions of the problem springs only (white) and after the settling period (red). c) Energy after the settling period versus the number of resets starting from random initial conditions before deformation (blue) and as deformation progresses (red).

In the following experiments we follow the previous protocol of periodic disturbances and to this we add an additional long settling period at the end of the experiment, Fig.5.b. During this final period the effective natural length of the L-springs converges to the distance between pairs of masses at the equilibrium of P-springs only, i.e., the L-springs exert no forces at equilibrium.

We examine two different kinds of external problems, specifically, a problem defined over continuous states (Scenario 2a), and a combinatorial optimisation problem defined over binary states (Scenario 2b).

#### Scenario 2a) Solving a general (continuous valued) external problem

We simulate *N* = 300 randomly distributed masses connected by P-springs 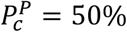 , with uniform distributed natural length 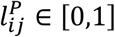 and uniform spring constants, *k*^*P*^ = 1. To this we add densely connected L-springs (connectivity 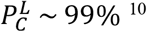 with uniform natural length 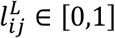 and uniform, and much weaker, spring constants 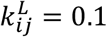 .

We plot the energy of the final equilibria of the learned system versus the number of disturbances Fig.5.c&d (red) and, for reference, plot this on a background of the energy of equilibria found when only P-springs are present (blue). Again, we find the system converges to low energy configurations of the P-springs. This is confirmed by a histogram of the energy of equilibria found from P-springs only Fig.5.c (blue), and P-springs plus final L-springs Fig.5.c (red). Again, to quantity the robustness we calculate the number of STDs between the final energy found and the mean energy of the problem springs over 10 independent realisations. Again, we found 80% of runs were over 3 STD from the mean and the average number of 3.0, indicating that the found solutions of this quality were extremely rare without the influence of the deforming springs.

#### Scenario 2b) Solving a discrete combinatorial optimisation problem

The previous examples demonstrated that, under relatively broad assumptions, a mass-spring-damper system can exhibit adaptation in the strict sense of learning to solve a problem better with experience. In the previous cases, the problem space is a continuous state space. Can this kind of adaptation by natural induction solve the kind of combinatorial optimisation problems that are more familiar in computer science? Combinatorial optimisation problems are a very general class, including MAX-SAT, TSP, etc.. The main departure from Scenario 2a is that the problem space in Scenario 2b is binary not continuous. For this we need to constrain the initial system architecture further to approximate binary states. This requires some additional mechanistic constraints (see Appendix 3 for details of Scenario 2b), but the natural induction mechanism is the same as before (Scenario 2a), i.e. using constant problem springs and a separate uniform network of deformable learning springs. We use the problem springs to define a spin glass system. Finding the lowest energy configuration of this systems corresponds to solving MaxCut graph partition problem and is NP-hard in general (Karp 2010).

As before, we find that adaptation by natural induction finds good quality solutions that are very rarely found without learning. We ran 10 runs and found that in 8 of these the binarized solution energy in the presence of learning was over 4 STDs away from the mean found without learning, demonstrating a significant increase in the quality of solutions found.

## 4. Discussion

### 1.9. The relationship between natural induction and natural selection

Adaptation by natural induction and adaptation by natural selection share a number of features: Both involve the incremental accumulation of small changes over time, both can result in the increased fit of an adaptive system to a system of constraints, and both involve simple gradient-following principles. They are also different in the mechanisms that they require (their necessary and sufficient conditions), how they work (their algorithmic principles) and, consequently, in their adaptive competencies. Whereas natural selection depends on the differential survival and/or reproduction of entities, natural induction operates by the differential easing of frustrated relationships between entities^11^. Moreover, the raison d’etre of selection as a theory of biological evolution is to avoid dependence on variation that is directed toward adaptive outcomes^12^, whereas natural induction exploits the fact that variation directed toward easing frustrated interactions is normal in physical systems and adaptively significant. These mechanistic differences are important, but it is perhaps the difference in their algorithmic principles and adaptive competencies that is more important. Even though both include gradient-following principles, they are different algorithms (Watson 2012), and this is clear because they have different problem-solving capabilities.

A common interpretation of adaptation by natural selection, i.e. characterised as a population following fitness gradients to a local peak in a fitness landscape (Wright 1932, Provine 1989), aligns well with local gradient-following principles of (first order) physical optimisation^13^. At a suitable level of abstraction both can be described as processes that follow local gradients to a local optimum. Clearly, though, they are mechanistically different ways of implementing this process. Specifically, natural selection can be described as a process of random generation and selective retention, whereas in the Newtonian model of a physical system, a ball, for example, is deterministically caused to move in a directional fashion by the reaction to the slope, i.e. to roll downhill not uphill. No population of balls, random variation or selection process need be involved. It is also possible, however, to think about a statistical mechanics process where the position of the ball is represented by a probability distribution of possible future positions, which is updated to amplify positions that are lower in energy compared to those that are higher in energy. So, does it really matter whether it is a statistical-mechanics (variation and selection) process or a Newtonian (directed learning) process? If the gradient-following outcome is the same, does the mechanism matter?

In some contexts, the mechanism seems to matter a lot to biological thought; Evolution by natural selection can only occur if there is a population, suitable variation, and selection – anything else, is not natural selection. On the other hand, it is common to equate evolutionary adaptation with a hill-climbing process, and to the extent that the change in an evolving population is non-arbitrary, this level of functional equivalence seems to capture its adaptive competence. However, if that is true, it has a curious implication; Although it is traditional for evolutionary thought to consider processes that go uphill (in a fitness landscape), whereas physical models go downhill (in an energy function), this does not make either one cleverer, i.e. a more effective optimiser, than the other (regardless of whether they are implemented in a statistical manner or a Newtonian manner). That would suggest that a literal ball rolling down a literal hill would constitute adaptation also. Is local hill-climbing sufficient to produce the biological adaptation we observe? Is the genius of Darwin’s theory just that it provides a hill-climbing process capable of operating in the appropriate organismic ‘design space’? Or do the mechanistic details of evolution by natural selection, or the context in which it occurs, matter to its adaptive competence (Watson 2012)? It seems likely that the details do matter, but to the extent that evolution by natural selection is characterised as a simple (substrate-independent) hill-climbing process it fails to capture these potentially important aspects.

The models presented here can be interpreted as literally physical systems, but they can also be interpreted as models that stand-in for other natural optimisation processes, including those involving the local gradient-following capacity of evolution by natural selection. Adaptation by natural induction is instantiated as a Newtonian process in the models we have illustrated – with forces and (directional) reactions rather than a statistical mechanics process (random variation and selection). And because they are different mechanisms this means that natural induction has different necessary and sufficient conditions to natural selection, and may apply in cases where natural selection does not (Watson and Lewens 2024). But in terms of adaptive competence, that is not the important difference between natural induction and natural selection. The problem-solving competence of natural induction is not the same as that of natural selection because they are different algorithms (not because they are different mechanisms for implementing the same algorithm). Adaptation by natural induction finds better solutions than a local gradient-following optimisation process, a.k.a. a hill-climber. If biological evolution is algorithmically equivalent to a hill-climber (first-order optimisation only), then its adaptive competence is inferior to adaptation by natural induction. Conversely, if biological evolution is a more sophisticated optimiser than a hill-climber, then the substrate-independent algorithm of random variation and selection does not describe it.

It might be appropriate to conceive biological evolution as a simple gradient process when natural selection acts on a simple vector of genes or vector of phenotypic traits individually determined by corresponding genes in a one-to-one fashion (Kounios, Clune et al. 2016). In this case, the action of evolution by natural selection is analogous to the bed of clay – a univariate model. That is, each selective coefficient is responsible for the change in frequencies of alleles at one locus. Other work has illustrated, however, that an evolutionary process operating on heritable variation in the connections of a dynamical network can exhibit the same kind of adaptive competence as natural induction. This includes gene-regulation networks (Watson, Buckley et al. 2010, Watson, Wagner et al. 2014, Kounios, Clune et al. 2016, Kouvaris, Clune et al. 2017, Brun-Usan, Thies and Watson 2020). In this case, the gene network constitutes a dynamical developmental process that generates phenotypes indirectly (and ‘disturbances’ are provided by a lifecycle that resets developmental states to a neutral, undifferentiated, epigenetic state). This gives it the possibility of representing relational interactions between traits (i.e. pleiotropic interactions in the genotype-phenotype map of such a developmental process may constitute an associative model of selected phenotypes (Watson and Szathmary 2016)). Previous work shows that under these circumstances the outcome is superior to local gradient optimisation (Kounios, Clune et al. 2016). That is, evolution by natural selection acting on the parameters of a developmental process (second-order) can solve problems that evolution by natural selection acting on a directly encoded phenotype (first-order) cannot.

One interpretation of this is that natural selection can exhibit the same adaptive competence as natural induction after all. However, adaptation by natural induction is not the algorithm that Darwin described (evolution by natural selection describe a process that provides model induction, a network of viscoelastic connections, an ability to learn to adapt better with experience, nor the significance of ‘pulse’ or disturbances, amongst other things). So, even though the basic gradient-following process might, in biological cases, be provided by a variation and selection process, it might be more correct to attribute the adaptive competence to natural induction, and not to natural selection. After all, whereas the theory of evolution by natural selection focusses on the mechanism of variation and selection, this paper demonstrates (by using purely Newtonian processes) that this superior adaptive competence arises from the dynamical feedback between model induction and optimisation, and not from random variation and selection. Whereas natural selection depends on the differential survival and reproduction of things, natural induction fundamentally depends on the differential easing of frustrated relationships between things. Although adaptation by natural induction is therefore fully compatible with a Darwinian model of evolutionary change, these are different adaptive algorithms with different necessary conditions, different algorithmic principles and different adaptive competencies.

We also note that the relationship between evolution and learning has been recognised and developed by many. At a suitable level of abstraction, evolution and some learning methods appear to be the same algorithm (i.e. ‘trial and error’ plus reinforcement equates to random variation and selection). Both can be understood as processes that optimise a function by, to a first approximation, following local gradients. It has often been noted that reinforcement learning and evolution by natural selection are closely analogous (Skinner 1981, Watson and Szathmary 2016), and indeed, the replicator equation (an abstraction of biological evolution under natural selection) and Bayesian updating (a learning optimisation process) have been shown to be formally equivalent (Harper 2009, Shalizi 2009). See also (Campbell 1983, Frank 2009, Valiant 2013, Chastain, Livnat et al. 2014, Kouvaris, Clune et al. 2017, Vanchurin, Wolf et al. 2021) for the relationship between learning and evolution. These works expand and deepen our understanding of the adaptation provided by natural selection. However, note that adaptation by natural induction involves a two-way feedback between an optimisation process and an inductive learning process – the latter on its own is simply an optimisation process in model parameters, and not sufficient to demonstrate an improvement in problem-solving competency.

The results in this paper thus demonstrate that a dynamical system described by a network of viscoelastic connections, and subject to occasional disturbances, exhibits adaptation in the more stringent sense of learning to optimise better with experience or improving its problem-solving competency over time – and this is not the same mechanism, algorithm or competency as natural selection.

Where might networks of suitable connections occur naturally? By far the most common examples of viscoelastic networks are in fact biological ones. Ecological networks, protein networks, cytoskeletal networks, metabolic networks, bio-electrical networks, social networks and the biosphere as a whole are all networks at least partially characterised by linkages that are likely to give-way under stress and are subject to at least occasional perturbations. Although these networks all involve biological individuals and materials, most of them are not (always) evolutionary units so natural selection does not straightforwardly apply. This suggests that the interaction of natural selection and natural induction may be complex and possibly widespread (Watson and Lewens 2024). Outside of systems that we already recognise as biological, another obvious candidate where natural induction may be important is the origins of life and origins of evolution (Pross 2004). To the extent that a pre-biotic chemical network has internal conformation structure that gives-way under stress we speculate that it has potential to induce a model of its past experience that can anticipate and generalise, without having properties sufficient to be a bone fide evolutionary unit.

Whilst we wish to make the case that the conditions for this are quite natural and not onerous (i.e. do not require selection or design), we do not claim that these conditions are ubiquitous or even frequently or commonly met. Bear in mind that the conditions for evolution by natural selection, namely self-replicating systems with heritable variation in reproductive success, are hardly ubiquitous in the physical world (even though they may be realised in various substrates (Hodgson 2005, Campbell 2011)) and their origin is not known. Instead, we claim that the contrary assumption, that natural selection is the only possible naturally occurring mechanism of spontaneous adaptation, is not correct. Who would make such a claim? Actually, this assumption is quite widely adopted, usually implicitly, with very wide reaching and important implications. We have developed the implications of this elsewhere (Watson and Lewens 2024).

### 1.10. Limitations

Natural induction does not occur unless particular conditions are met. It requires a set of state variables and a set of structural parameters such as a network of connections (interaction terms in the dynamical system), these connections need to give-way slightly under stress, and the system needs to be subject to disturbances (or episodic stress). These conditions are not uncommon in natural networks but, of course, not universal. We have made a number of specific assumptions in the particular systems we have illustrated (spring constants, timing parameters, connectivity, etc.), but the central claim of this paper is robust to these choices and other details; that is, natural selection is not the only possible source of spontaneous adaptation. The aim of our illustrations is not to be the ‘last word’ or definitive case on such issues, but merely to open-up the discussion and fuel debate that is productive.

Nonetheless, in order for natural induction to produce adaptation (i.e. increase in optimisation capability) some general conditions are required. These correspond to the conditions for good generalisation in a learning system. The problems solved here are constructed from pairwise constraints. These capture a large class of problems, but not all. It is of course, easy to construct optimisation problems that natural induction cannot solve. The state dynamics need to spend most of their time at configurations that are better than average (better than random). This condition is easily met and the condition that they spend most of their time at local optima, as in our illustrations, is probably not necessary but not yet investigated. Likewise, we suspect that it is not essential that interactions are symmetric (which guarantees only fixed point attractors) but asymmetric interactions and non-fixed point dynamics are not investigated in this paper. Additionally, the system must be subject to disturbances – not too frequent to prevent the system from spending most of its time at good configurations, but not too infrequent that it does not visit a representative sample of good configurations. Disturbances do not necessarily need to be complete resets of the state configuration in order for some induction to occur, but we anticipate that partial resets will limit the independence of the samples from which associations are being learned. This is a complicated matter however because partial resets to state variables can sometimes act in a similar manner to updates in interaction parameters. Together, these conditions correspond to the fact that learning works well only when the training data is representative of the class that must be learned – you cannot learn a general class from a single example (or an impoverished distribution of samples) and that is why the disturbances are needed. Learning rate also matters (particularly to online learning where there is positive feedback between what is learned and the data that is learned from) because you do not want learning to take unnecessarily long nor for the system to converge on the first (arbitrary) state it experiences.

Finally, natural induction is an inductive learning process and there is an important fundamental limitation to any inductive learning process – the need for a suitable *inductive bias*. Generalisation cannot occur without induction, but any general rule is necessarily under-determined by past experience. Over the set of all possible general rules that are compatible with the data, any prediction is possible (e.g. “all swans are white except the next one, which is pink”). Therefore, any inductive learning process can only produce a prediction (right or wrong) by preferring particular generalisations over others. This preference is not determined by the data (by definition) and is known as inductive bias. An inductive bias can be as simple as a preference for simple models over complex models (a.k.a. parsimony pressure, or regularisation in machine learning).

The simplest kind of model capable of representing associations, and thus capable of non-trivial generalisation, is a correlation model; and this is the inductive bias underlying our results. That is, in the examples presented here (and others (Watson, Wagner et al. 2014, Power, Watson et al. 2015, Kounios, Clune et al. 2016, Watson and Szathmary 2016, Kouvaris, Clune et al. 2017)), the model space in which induction occurs is built from pairwise interactions – like neural networks are. This works well in many learning problems because it is as simple as possible but not more so (generalisation with a univariate model, like the bed of clay, is limited to simple similarity measures (Watson and Szathmary 2016)). For our worked examples above (and in the previous work), correlation learning is a good inductive bias because the constraints that determine the structure of the problem are also built from pairwise interactions. This is why generalising over a distribution of some particular local optima is able to predict the location of (i.e. enlarge the attractor for) other local optima that have not been previously visited. In general terms, the implicit inductive bias will be suitable whenever the model and the problem are built from a similar causal geometry (in this case, a network of pairwise interactions) – this is natural when a system is learning by adjusting its own connections (Watson, Buckley and Mills 2011).

The acknowledgement of an inductive bias is not to suggest in any way that the model was somehow given the solution in advance. All learning requires induction, and a learning process does not know answers in advance, it acquires this information from experience. Although it is not usually presented as such, all optimisation is really a task that requires induction. That is, an optimisation process must predict the location of (hard-to-find) good solutions from (easy-to-find) samples. If, conversely, an adaptive process had already visited the location of good solutions, then the problem is already solved. Even a simple hill-climbing process, or an evolutionary process, is no better than random guessing if it is not employing a suitable inductive bias – in this case, the assumption of local smoothness (Wolpert and Macready 1997). To visualise this, imagine a natural selection process on a truly random fitness landscape with no auto-correlation – here the location of any solution sampled in the past provides no information whatsoever about the location of potentially better solutions, and thus natural selection has no optimisation ability.

In adaptation by natural induction, we are simply exploiting a similar auto-correlation bias but in a slightly deeper representation instead of the original features. Natural selection depends on the assumption that good solutions have state *values* that are similar to the values found in other good solutions. In natural induction, the implicit assumption is that good solutions have *correlations* among their state values that are similar to the correlations found in other good solutions. In difficult problems, the simple value-based assumption is limited and the correlation-based assumption provides a little more competence. Problems that are even more difficult, if they have any learnable structure at all, require deeper models and the principles exploited by natural induction can be extended in this direction (Caldwell, Knowles et al. 2021). Recognising that adaptation requires learning, and learning requires generalisation, and generalisation requires an inductive bias, helps us to understand how adaptation really works and what is required. Without an inductive bias all adaptation (including natural selection) would be magical.

We have not yet investigated systems with hidden state. Although the problem springs can represent a problem that is external to the learning springs, in the examples illustrated thus far all the state variables are shared. Inducing a model of a complex system that has hidden state can be much more difficult, and reflexively, a learned model that contains hidden state (or a ‘deep’ representation) can express relationships that a shallow model cannot (Caldwell, Watson et al. 2018, Caldwell, Knowles et al. 2021, Watson and Levin 2023).

In the particular system of masses and springs used here, where the parameters of the problem and the induced model are embodied in spring lengths, we show that the change in the parameters is given by the differential of the potential energy function, see Appendix 1. This seems intuitively natural – forcing the state variables causes the internal structure to give-way in the direction that reduces the stress in the system and lowers the potential energy of that state. This also enlarges the dynamical attractor of that state configuration. The generality of this concept for other physical systems and other structural variables is not known but some other possibilities have been demonstrated. For example, earlier work modelled changes in interaction strengths analogous to spring constants rather than a spring lengths (Watson, Buckley and Mills 2011, Watson, Mills and Buckley 2011, Watson, Wagner et al. 2014, Power, Watson et al. 2015)^14^.

The illustrations of natural induction presented here using masses and springs have all been conducted in a heavily-damped regime which minimises oscillatory. We have not yet investigated the kind of adaptive algorithm that is instantiated in the oscillating regime. This is obviously a lot more complicated but we suspect that it might be quite interesting (Wang and Roychowdhury 2019). There are possibilities of states being represented by phases, interactions being represented by resonance properties, and learning represented by harmonic phase locking that is natural between coupled oscillators. Preliminary work in another context suggests that phase synchronisation could play an important role in scaling-up the adaptive process from one level of organisation to another (Watson 2023, Watson, Levin et al. 2023).

## 5. Conclusions

It has been argued strongly, and widely assumed across biological thinking, that natural selection is the only possible mechanism capable of producing spontaneous adaptation in natural systems. Here we show that this assumption is false; adaptive optimisation occurs spontaneously in physical systems with suitable natural properties, through an effect we call adaptation by natural induction. This occurs in dynamical systems described by a network of viscoelastic connections subject to disturbances. A viscoelastic connection is simply one that ‘gives way’ slightly under stress, which is a natural property of many physical materials, biological networks, and complex systems more generally. When disturbances cause the system to visit a distribution of locally-optimal solutions, the changes to the connections in the network learn a generalised associative model of the solutions visited, which causes the system to adapt in the rigorous sense of improving its problem-solving competency. The simplicity of natural induction, and its necessary and sufficient conditions, offers a solution to both the chicken and the egg problems – i.e. natural selection is not involved at run time nor in the construction or setup of the system. This has important implications for our understanding of biological evolution, and adaptation in other complex systems – not least that adaptation can occur spontaneously in systems that are not units of selection (Watson and Lewens 2024).

## Acknowledgements

The authors thank Chrisantha Fernando, Adam Davies, Dave Prosser, Freddy Nash, Jamie Caldwell, Tobias Uller, Kostas Kouvaris, Christoph Thies, Tazzio Tissot, Jonathon Young, Lucas Mathieu, Samuel Lennard, and Joshua Knowles for discussion. The authors gratefully acknowledge the support of grants 62230 & 62220 (RW) and 62212 (ML), from the John Templeton Foundation, and BBRSC grant BB/P022197/1 and the UKRI Horizon Europe Guarantee scheme as part of the METATOOL project (CB).

## Appendix 1

**Appendix Figure 2:**
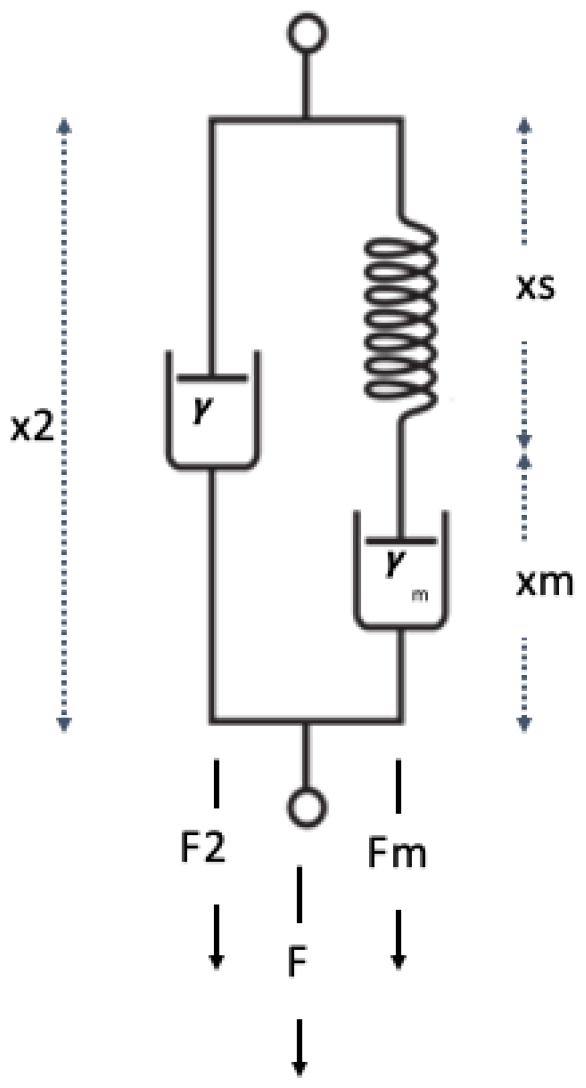
A maxwell configuration comprising of a spring in series with damper, in series with a second damper.

Here we derive the equations of motion for the Maxwell spring by resolving forces in the normal way. First, we define the displacement of each element thus,

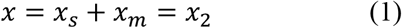

where *x* is the length of the entire spring and and *x*_*m*_ and *x*_2_ are the lengths of the ideal spring, damper in series and in parallel respectively, see App. Fig. 1.

The force on the unit mass is given by,

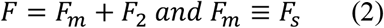

where *F* is the total force and *F*_*s*_, *F*_*m*_ and *F*_2_ are forces in the ideal spring, damper in series and in parallel, respectively. Writing,

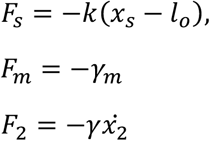

We can we replace 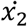 with 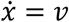 because we can see from Eqn. 1 this is just the velocity of the combined system. Taking the time derivative of Eqn. 1 and substituting in Eqn. 2 we arrive at,

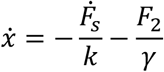

Rearranging this equation and using Newton’s law yields the equations of motion for the unit mass as,

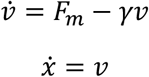

To convince ourselves this is correct consider the case where the dashpot is stiff and effectively rigid, i.e. , *γ*_*m*_ → ∞. Integrating, 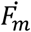 and taking initial conditions we return a simple damped spring as,

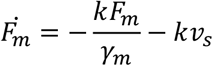

We can show this is equivalent to the ‘learning’ rule in Eqn. 1 in the main text by again integrating 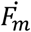

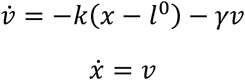

where we have made the substitution,

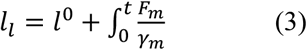

Now differentiating both sides of Eqn. (3) and substituting *F*_*m*_ with the force on the spring, i.e.,*F*_*m*_ = −*k*(*x* − *l*_*l*_) we get

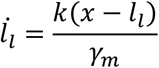

with learning rate 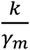 .

Another way to arrive at this learning rule is to take the derivative of the energy with respect to the parameter. Specifically, consider the energy of the above system,

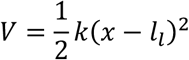

And defining update that minimises this energy by gradient descent.

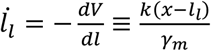

## Appendix 2

**Appendix Figure 2:**
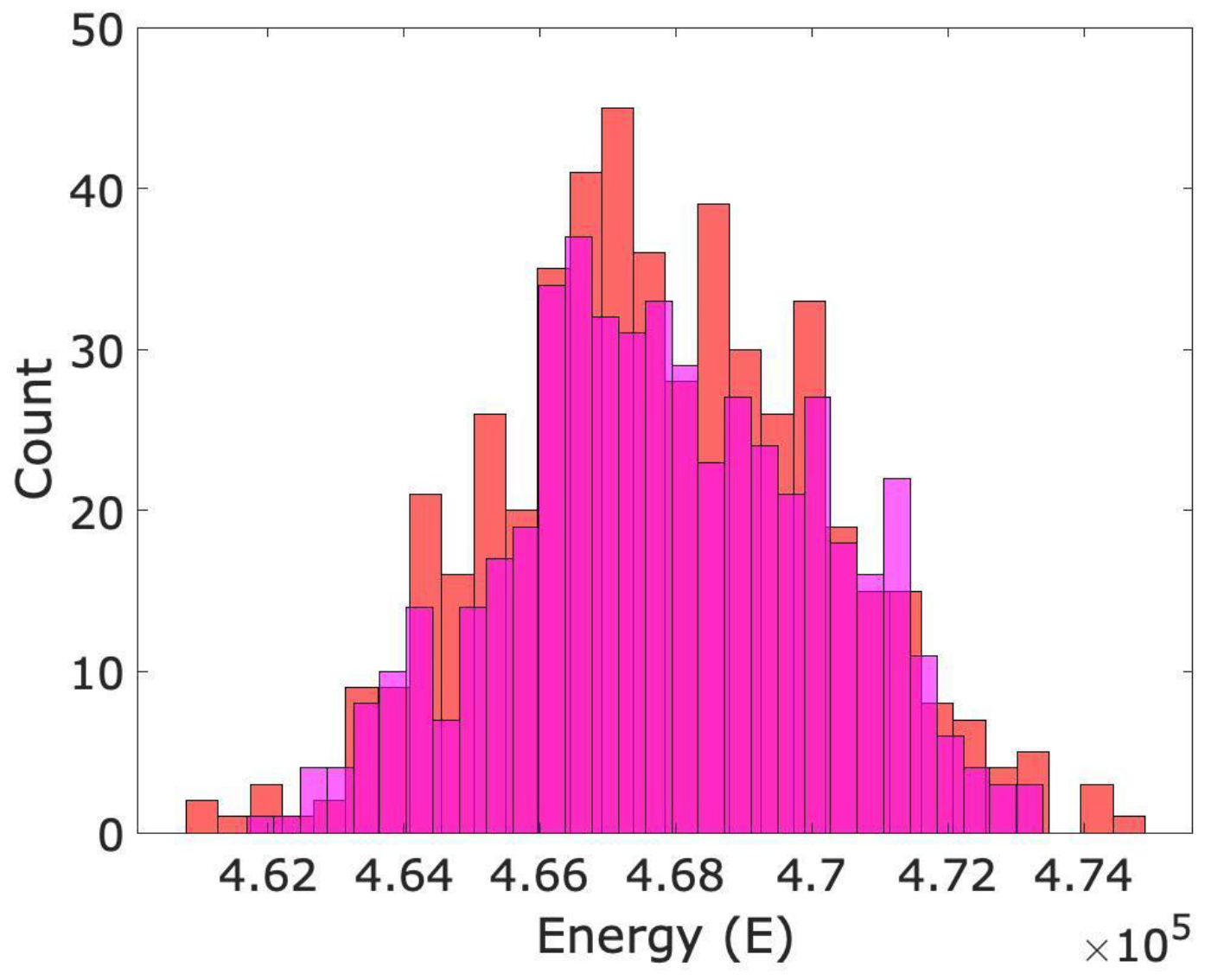
The distribution of found energies on the P-spring alone (red) verses running them with P- and L-springs (before learning) but then removing the L-springs (magenta). The distributions are almost identical. We can thus think of L-springs as the addition of an initially neutral field of plastic material that (initially!) does not alter the problem.

## Appendix 3

In Scenario 2b, we examine a combinatorial optimisation problem specified as finding the minimal energy state of a spin glass system. We consider network of *N* binary spins *s*_*i*_ ∈ {−1,1} with symmetric interactions, *J*_*ij*_ = *J*_*ji*_ , and *J*_*ii*_ = 0, but randomly connected *J*_*ij*_ *∼ U*{0, −1} with energy defined by *H* = − *∑*_*ij*_ *J*_*ij*_*s*_*i*_*s*_*J*_.

Finding the lowest energy configuration of this systems corresponds to solving MaxCut graph partition problem and is NP-hard in general (Karp 2010). To represent this problem in our mass-spring-damper system we consider the following setup. Each mass is attached to a vertical rod constrained to travel vertically and tied with a spring to a fixed point at height *y* = 0 (Fig.A3.1.a), which effects bistability (i.e. the binarisation) in the y position. Pairs of masses are randomly connected with a set of P-springs as before. To simplify our simulation we ignore spatial constraints and assume that every pair of rods is a unit length apart. Resolving forces we can write the potential energy of this systems as,

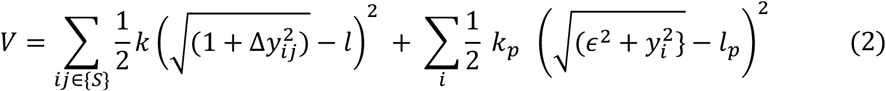

where *y*_*i*_ is the vertical displacement of each mass with respect to the fixed point, 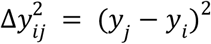 is the y-distance between masses, and γ is damping effected by friction on the rod. The natural length and spring constant of the P-springs and tie spring are *k, l* and *k*_*p*_, *l*_*p*_respectively. The distance of the tie to rod is given by ϵ. The first term of this equation is the potential energy due to the P-springs, note by construction there are no net forces in the x-direction. The second term is the contribution to energy of the tie spring. In the absence of P-springs, and with the tie point close to the rod (ϵ → 0) the system is driven into either up or down position, *y*_*i*_ = ±*l*_*p*_. This bistability is preserved in the presence of P-springs if *k*_*p*_ > *k* such that the second term in Eq.2 dominates the first. Adding P-springs which are much longer than unit length (i.e. longer that the distance between pairs of rods) forces coupled masses to misalign and emulate the behaviour of a negative weight in a binary spin glass system. We can see this by noting that when the natural length of the P-springs is much larger than the separation of rods, *l*^0^ ≫ 1, the first term in LHS of the energy Eq.2 is dominated by −2*L*|*yi* − *y*_*j*_| and the masses separate^15^.

As before, *N* = 300 with connectivity 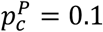 and parameter *k* = 1, *l* = 10, *k*_*p*_ = 1, *l*_*P*_ = 1, *γ*. The equivalent spin glass problem can be constructed by translating all long springs in {*S*} into negative weights in *J*_*ij*_ = −1. We run the mass-spring-damper to equilibrium and interpret these as solutions to the spin glass problem by interpreting all rods above and below the mid-point of travel as negative and positive spins respectively (Fig.A3.2.b). We compare the performance of this system on the MaxCut problem against standard hillclimbing algorithm (HC) where we flip each spin with probability P=1/N and run it for 20000 steps. In Figure A3.1.c we see that the distribution of solutions found (before L-springs deform) are comparable to the solutions found by a hillclimbing algorithm.

To allow the system to adapt, as before, we introduce a highly connected set of L-springs (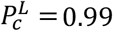 and *k*^*L*^ = 0.1 and *l*^*L*^ = 20). We follow the procedure as in Scenario 2 with periodic disturbances, and after some number of disturbances we then let the system relax for a long period to read out the found solution. In Fig.A3.c we plot the energy of the found solutions against the number of disturbances (red dots) as well as the energies found with P-springs alone (blue dots) demonstrating again the system finds better solutions over time. In Fig.A3.d we convert the binarised solution states into their equivalent MaxCut solution. We ran 10 runs and found that 8 of these the binarized solution energy in the presence of learning was over 4 STDs away from the mean found without learning, again demonstrating a significant increase in the quality of solutions found.

**Appendix Figure 3.1:**
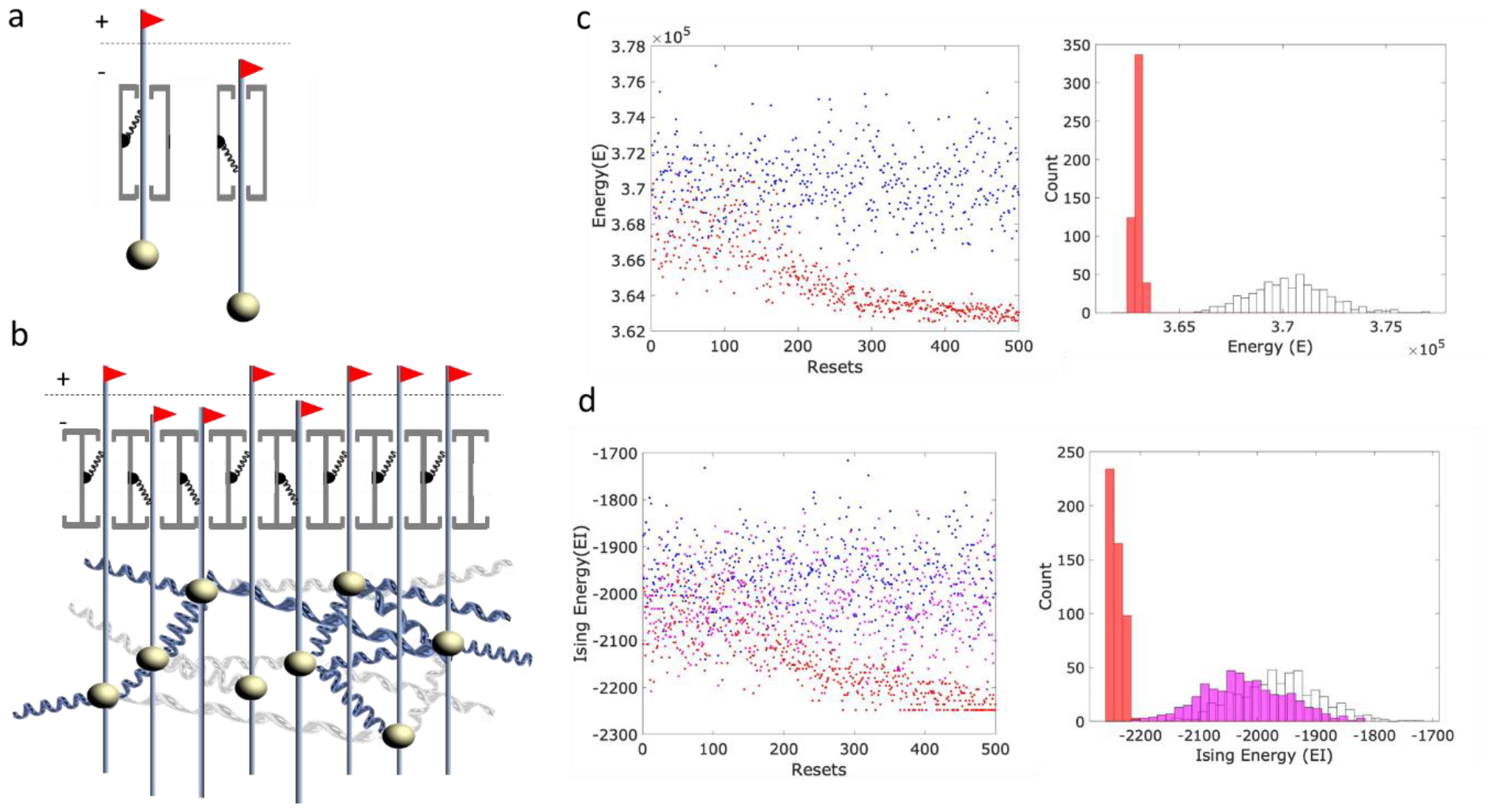
Adaptation by natural induction discovers solutions to a binary constraint problem (Scenario 3). a) A system of masses set on rods constrained to move in the y-direction. Each rod is tied to the y-axes with a short spring which, in the absence of other springs yields a bistable resting position in an up/down position. b) Pairs of masses are also connected by a set of elastic long springs (‘problem springs’, blue). These springs are longer than the distance between the rods in the horizontal plane and they thus have the effect that the masses want to misalign. Masses are also densely connected with set of uniform-length deforming spring (‘learning springs’, grey). A binary readout of the displacement of the masses is interpreted based on whether the tie spring is oriented upward or downwards (i.e. red flags above/below midpoint). c) Left: The energy of the final configuration of springs verses the number of resets, without learning springs (blue) and as learning springs deform over time (red). Right: a histogram of the energy of the equilibria found for just the problem springs (white bins) and from different initial conditions for the final L-spring lengths (red bins). The L-springs allow the system to consistently visit low energy solutions defined by P-springs. Furthermore, the lowest energy state was extremely unlikely to have been visited without the deformed L-springs. d) Same as c) but here the final state is binarized and energy measured on the underlying Ising energy of the problem. Again, red and white bins are with and without L-springs, respectively. The magenta bins show the distributions of solutions from multiple runs of the hillclimber.

**Appendix Figure 3.2:**
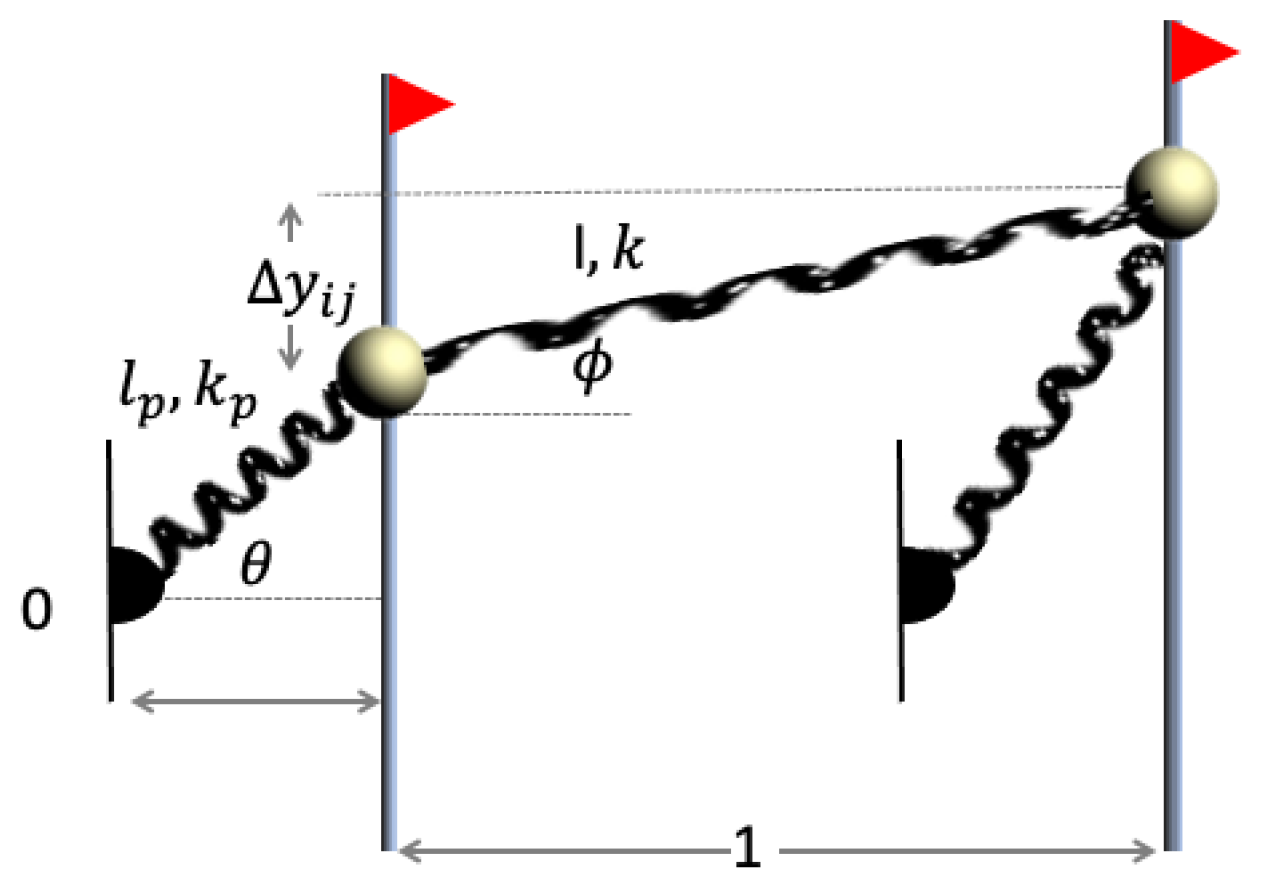
Two masses constrained by metal rods to move in the y-direction. *The rods are tied to a pivot point by a tie spring*.

Resolving forces in the y-direction for the tie spring we have,

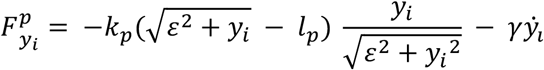

Where we have used *sin* 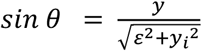 and we assume some friction on the vertical rod. Simplifying, we get,

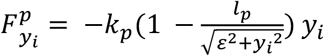

Similarly, resolving forces for the spring between masses i and I we have

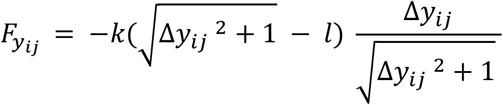

Where Δ*y*_*ij*_ = *y*_*j*_ − *y*_*j*_ and which simplifies to,

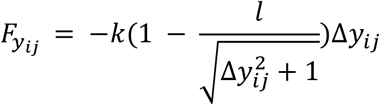

This the total forces on mass *y*_*i*_ is

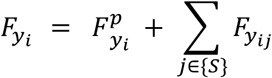

And the potential energy of this system,

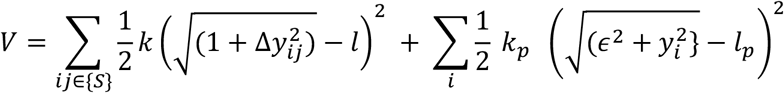

## Footnotes

1 Whilst there is a long history exploring the idea that cognitive processes can indeed be characterised as variation and selection processes (Campbell 1960, 1983, Edelman 1987, Fernando et al. 2010, 2012), artificial neural networks show that this type of learning can be implemented with gradient methods that do not involve variation and selection. Gradient methods do not involve selection at any level of organisation including the level of synapses, neurons nor at the level of the network as a whole.

2 i.e. modelling each variable independently

3 Hebb’s rule (Hebb 1949), a simple method of updating neural connections often described as ‘neurons that fire together wire together’, is equivalent to energy minimisation on the weights of the network given the current (e.g. forced) pattern of the states (Watson et al 2010, 2011a, 2011b, 2014).

4 We are not intending to advance or augment a general understanding of induction or inductive inference in this paper; ‘natural induction’ merely aims to acknowledge the importance of inferences that are under-determined by observations, and their necessity for generalisation and adaptation, in the effect we are modelling.

5 Note that evolution by natural selection is often characterised in this way, i.e. as independent selective coefficients acting on a vector of independent alleles, or non-pleiotropic traits. In this case, selection is not capable of inducing a generalised model of past selective conditions. However, when selection acts of the parameters of a developmental process, with complex pleiotropic interactions, it is possible to store and recall multiple fit phenotypes in a single genotype and for generalisation in this model space to produce novel phenotypes from the same class (Watson et al 2014, Kouvaris et al 2017).

6 A less value-laden conception of evolution as a dynamical process, without a problem to be solved, and without a pre-existing niche to be occupied, is acknowledged (Gould & Lewontin 1979, Levins & Lewontin 1985). However, if adaptation is construed as merely ‘whatever happens as a result of natural selection’, for example, then it remains tied to natural selection. Adaptation needs to address a design-like property (independent of natural selection), and problem solving is one way to characterise this. At the least, a process that can provide a non-trivial problem-solving competency is a high bar for assessing adaptation.

7 There is some possibility that ‘abduction’ or ‘transduction’ are more accurate terms but we feel that induction captures the basic flavour sufficiently.

8 The extent to which this results in effective adaptation (optimisation better than a local hill-climber) will depend on the suitability of the inductive bias implicit in the system architecture, i.e., whether the causal geometry of the system doing the learning is like that of the environment it is learning about – See Discussion.

9 The quality of a solution therefore cannot be directly read off from the final state configuration in Scenario 1 because the original spring parameter values that represent the problem are lost as the springs change. This is why it was necessary to take each of the new configurations it finds and assess their quality by finding the nearest attractor in the original system (as described above).

10 We used 99% to avoid instabilities caused by symmetries of fully connected systems.

11 The accommodation of internal connections to a state configuration has some similarity to the over-production of neural connections and their differential retention or reinforcement by ‘selective stabilization’ which may also results in increased response to or memory of activation patterns Changeux, J.-P. and A. Danchin (1976). “Selective stabilisation of developing synapses as a mechanism for the specification of neuronal networks.” <u>Nature</u> **264**(5588): 705-712.. However, natural induction is a physical model without any population or selection process whereas selective stabilisation depends on an over-produced population of connections, and in terms of algorithmic competence, natural induction demonstrates an increase in adaptative capabilities. // The emphasis of natural induction on the differential easing of frustrated interactions agrees with an emphasis on networked integration ‘survival of the fitted’ (in contrast to ‘survival of the fittest’) Cohen, I. R. and A. Marron (2023). “Evolution is driven by natural autoencoding: reframing species, interaction codes, cooperation and sexual reproduction.” <u>Proceedings of the Royal Society B</u> **290**(1994): 20222409..

12 Darwin suggested that some variation was developmentally-environmentally directed, but was not specific about its adaptive significance. The nature of developmental bias and phenotypic plasticity, and the potential of these and other factors to influence genetic evolution adaptively, is an active topic Levin, M. (2023). “Darwin’s agential materials: evolutionary implications of multiscale competency in developmental biology.” <u>Cellular and Molecular Life Sciences</u> **80**(6): 142, Livnat, A. and A. C. Love (2024). “Mutation and evolution: Conceptual possibilities.” <u>BioEssays</u> **46**(2): 2300025..

13 It is acknowledged that biological evolution is not necessarily a good optimiser (Gould & Lewontin 1979), the idea of natural selection climbing gradients in a static fitness landscape is a serious over-simplification (Levins & Lewontin 1985), identifying any quantity that natural selection maximises is problematic (Grafen 2009), and the conception of natural selection as a problem-solving process has been criticised (Levins & Lewontin 1985). These are mostly arguments that weaken the interpretation of natural selection as an optimisation process, i.e. natural selection is, a least sometimes, less effective at optimisation than a local gradient process. Accordingly, doing better than local optimisation is a conservative criterion for adaptation.

14 If springs only weaken, and never increase in strength, this describes an update rule that is equivalent to one ‘half’ of Hebb’s rule (i.e. the half that *decreases* the magnitudes of weights). The limit of this is that ultimately all interactions (L-springs) go to zero and all states are independent of other states (except for influence of P-springs). This also depends on the differential reduction of frustrated correlation parameters (without any increases of correlation parameters that are not frustrated) so the initial effect of L-springs cannot be zero. Nonetheless, the current work likewise depends on differential easing and demonstrates that this is sufficient for significant adaptive problem solving (even when P-springs cannot be changed).

15 It would also be possible to emulate positive weights when the natural length the P-springs are smaller than the distance between rods, *l*^0^ ≪ 1

